# Structural basis of nucleosome recognition by the conserved Dsup and HMGN nucleosome-binding motif

**DOI:** 10.1101/2025.01.06.631586

**Authors:** Jaime Alegrio-Louro, Grisel Cruz-Becerra, James T. Kadonaga, Andres E. Leschziner

## Abstract

The tardigrade damage suppressor (Dsup) and vertebrate high mobility group N (HMGN) proteins bind specifically to nucleosomes via a conserved motif whose structure has not been experimentally determined. Here we used cryo-EM to show that both proteins bind to the nucleosome acidic patch via analogous arginine anchors with one molecule bound to each face of the nucleosome. We additionally employed the natural promoter-containing 5S rDNA sequence for structural analysis of the nucleosome. These structures of an ancient nucle-osome-binding motif suggest that there is an untapped realm of proteins with a related mode of binding to chromatin.

## Introduction

The specific interaction of proteins with the nucleosome is central to the regulation of chromatin function and dynamics. Our understanding of this aspect of chromatin regulation is limited, however, by our knowledge of the basis of nucleosome recognition via conserved protein motifs. In this study, we investigated the interaction of the high mobility group nucleosome-binding (HMGN) protein motif with the nucleosome. The canonical HMGN proteins are abundant factors that are present only in vertebrates, all of which have at least three *HMGN* genes (González-Romero et al. 2015; Nandury et al. 2020). The HMGN proteins bind to two high affinity sites on the nucleosome in a DNA-sequence-independent manner via a conserved RRSARLSA motif (Mardian et al. 1980; Sandeen et al. 1980; Ueda et al. 2008). The interaction of this motif with the nucleosome has been examined by NMR and modeling (Kato et al. 2011), but its high-resolution molecular structure has not yet been experimentally determined.

Intriguingly, an HMGN-like motif is present in the tardigrade-unique damage suppressor (Dsup) protein, which is a nucleosome-specific binding factor that has been shown to protect fly, plant, yeast, and human cells from DNA damage (Hashimoto et al. 2016; Chavez et al. 2019; Kirke et al. 2020; Ricci et al. 2021; Zarubin et al. 2023; Aguilar et al. 2023). The HMGN-like motif in Dsup is essential for binding to nucleosomes and for the protection of DNA from damage by hydroxyl radicals (Chavez et al. 2019). At present, Dsup is the only non-vertebrate protein that has been found to bind to nucleosomes via an HMGN-like motif.

To understand the relationship between the nucleo-some-binding motifs of Dsup and a canonical HMGN protein as well as to gain insights into their specific interactions with the nucleosome, we used cryo-electron microscopy (EM) to determine the structures of the Dsup-nucleosome complex and the HMGN2-nucleosome complex. In addition, to study nucleosome structure in the context of active chromatin, we analyzed nucleosomes that were reconstituted with a promoter-containing DNA sequence from a *Xenopus borealis* somatic *5S rDNA* gene (Rhodes 1985). These studies revealed that the 5S rDNA sequence is a useful alternative to the commonly employed synthetic or α-satellite DNA sequences (Armeev et al. 2022).

## Results and discussion

### Two Dsup molecules bind to the nucleosome

We initially sought to investigate the molecular interactions of Dsup with the nucleosome. To this end, we reconstituted the Dsup-nucleosome complex with recombinant *Ramazzottius varieornatus* Dsup (Fig. 1A) and the nucleosome core particle containing the 601 nucleosome positioning sequence (Lowary and Widom 1998) and performed cryo-EM analysis. Data collected from a single cryo-EM grid with a crosslinked sample resulted in two Dsup-bound nucleosome structures that mostly differ in the dynamics of the DNA ends. We determined the directionality of the DNA sequence in the cryo-EM maps using B-factor analysis (Supplemental Fig. S1; Supplemental Materials and methods). While the left outer gyre of the DNA is flexible in both structures, the right DNA end is fully wrapped in “Structure I” (2.7 Å resolution) but partially unwrapped in “Structure II” (2.8 Å resolution) (Supplemental Figs. S1, S2 and Supplemental Table S1). Importantly, the two structures contain extranucleosomal densities that correspond to two Dsup molecules bound to either side of the disk (Fig. 1B; Supplemental Fig. S2). We refer to Structural basis of nucleosome recognition by the conserved Dsup and HMGN nucleosome-binding motif the Dsup molecules as “A” and “B” depending on whether they are bound to the face of the histone octamer proximal to the left or right half of the 601 DNA, respectively. The highest local resolution of the Dsup densities is 3.1 Å (Supplemental Fig. S1), which allowed us to model residues 361 to 369 of both Dsup molecules (Fig. 1; Supplemental Fig. S3). Notably, this segment contains an arginine rich motif (RRSSR) that is part of the Dsup region that is similar to the core nucleosome-binding domain of the vertebrate HMGN proteins (Chavez et al. 2019). In addition, we obtained several 3D classes of Dsup-nucleosome complexes showing low resolution Dsup densities in different locations, suggesting that Dsup can transiently adopt multiple conformations around the nucleosome (Supplemental Fig. S4). Hence, two Dsup molecules can bind to the nucleosome through their HMGN-like motif.

**Figure 1.**
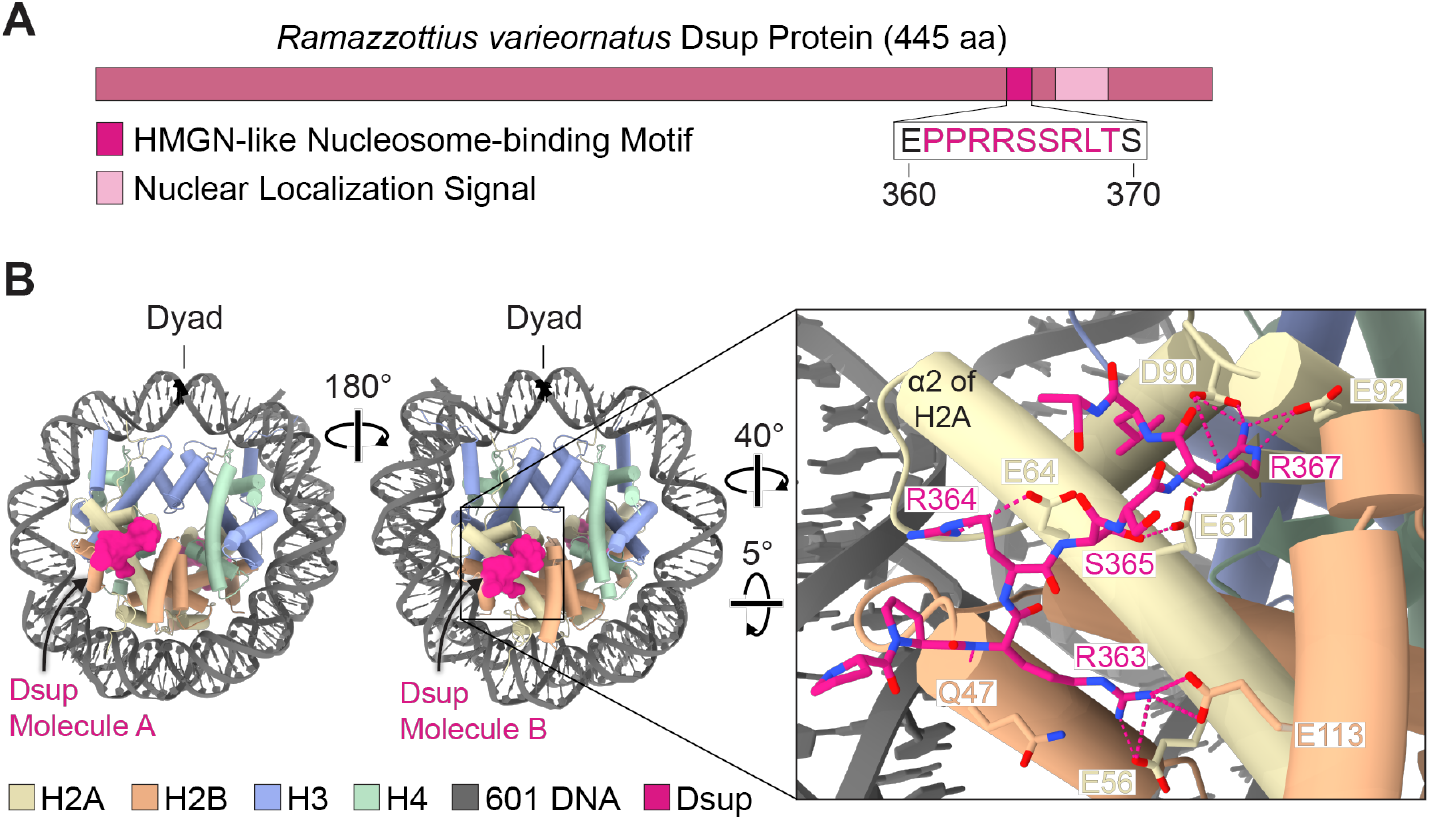
Two Dsup molecules bind to the nucleosome. (*A*) *R. varieornatus* Dsup protein. The HMGN-like nucleosome-binding motif of Dsup is highlighted (residues 360-370; Chavez et al. 2019). The sequence in pink type corresponds to the Dsup region modeled for each of the Dsup molecules in our structure. (*B*) Model of Structure I (with one wrapped end and one unwrapped end; Supplemental Fig. S2) for the Dsup-bound 147-bp 601 nucleosome crosslinked with glutaraldehyde. One Dsup molecule binds to each side of the disk. We refer to the two nucleosomebound Dsup molecules as Dsup molecule A and Dsup molecule B. The close-up on the right shows the interactions (dashed lines) of Dsup molecule B with the nucleosome acidic patch and with Q47 of the H2B α1-L1 elbow.

### Dsup interacts with the nucleosome acidic patch

Most of the interactions of the HMGN-like motif of Dsup with the nucleosome are shared between structures I and II (Fig. 1; Supplemental Fig. S3). In both structures, the PPRRSSRLT segment of Dsup straddles the long alpha helix (α2) in H2A and has multiple anchor points on the surface of the nucleosome acidic patch and the H2B α1-L1 elbow (Fig. 1B; Supplemental Fig. S3). Our analysis revealed that the backbone of Dsup R363 and H2B Q47 interact via hydrogen bonding, whereas the side chain of Dsup R363 forms salt bridges with H2A E56 and H2B E113 (Fig. 1B; Supplemental Fig. S3). Furthermore, Dsup R367 interacts with H2A E61, D90, and E92 (with the exception of Dsup molecule A in structure II, in which E92 is pointing out) and thus functions as the arginine anchor that is found in most acidic patch-binding proteins (Fig. 1B; Supplemental Fig. S3; Barbera et al. 2006; McGinty and Tan 2021). Dsup R364 interacts with H2A E64, but it was poorly resolved in one Dsup molecule, suggesting that it is more loosely attached than R363 and R367, which are buried within the acidic patch. In addition, Dsup S365 contributes to nucleosome binding through hydrogen bonding with either H2A E61 or H2A E64 (Supplemental Fig. S3). We also identified one class with Dsup density spanning the histone octamer (Supplemental Fig. S4A). In this class, the Dsup density extends from the N-terminal helix of H3 (H3 αN), the docking domain of H2A, and SHL +6.5 over the H2A-H2B dimer, and then to the vicinity of helix 1 of H3 and H3 loop L1 of the second histone H3 molecule. Overall, these findings indicate that the binding of Dsup to chromatin involves interactions with the acidic patch as well as with other regions of the histone octamer, and are consistent with a model in which the nucleosome-bound Dsup molecules function as a shield against DNA damage through their flexible regions (Chavez et al. 2019; Aguilar et al. 2023).

### Two HMGN2 molecules bind to the acidic patch regions in the nucleosome

We next expanded our cryo-EM studies to investigate the molecular interactions between a canonical HMGN protein and the nucleosome. To this end, we collected data from a crosslinked HMGN2-nucleosome sample containing human HMGN2 and a 5S rDNA nucleosome with 10 bp of linker DNA at each end. The 5S rDNA sequence contains the promoter region of a somatic *X. borealis 5S rDNA* gene. We obtained a 2.9 Å resolution map of the HMGN2-bound nucleosome (Supplemental Fig. S5 and Supplemental Table S1). The map shows density for one full wrap of DNA around the histone octamer but lacks signal for the 10 bp linkers and most of the outer turns (Supplemental Figs. S5, S6), which suggests high DNA flexibility. Two HMGN2 molecules, one on each face of the nucleosome at the acidic patch, are observed in the cryo-EM map of the HMGN2-nucleosome complex (Fig. 2B; Supplemental Fig. S6). The highest local resolution of the HMGN2 density on each side of the disk is 3.0 Å (Supplemental Fig. S5), including residues 22 to 26 of HMGN2, an arginine rich sequence (RRSAR) that is conserved in all HMGN proteins (Fig. 2; Supplemental Fig. S6; Ueda et al. 2008; Chavez et al. 2019).

**Figure 2.**
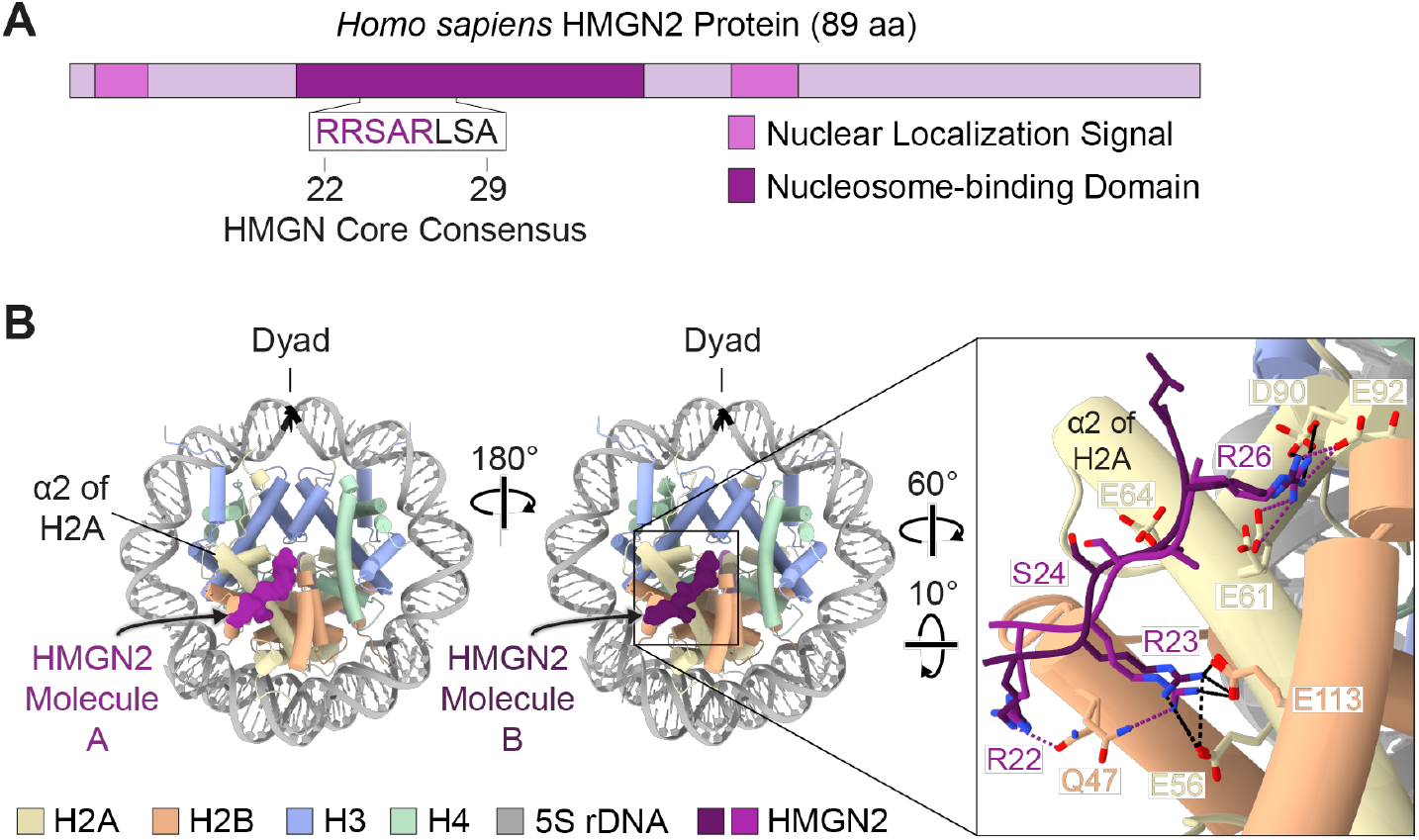
Two HMGN2 molecules bind to the nucleosome. (*A*) *H. sapiens* HMGN2 protein. The sequence corresponds to the core consensus motif that anchors the HMGN proteins to the nucleosome (HMGN2 residues 22-29; Ueda et al. 2008). The purple type highlights the HMGN2 region modeled for each of the two HMGN2 molecules in our structure. (*B*) Model of the HMGN2-bound 167-bp 5S rDNA nucleosome. We refer to the two nucleosome-bound HMGN2 molecules as HMGN2 molecule A and HMGN2 molecule B. The binding of the two HMGN2 molecules to the nucleosome is not identical. The close-up on the right shows the shared (black dashed lines) and distinct (purple dashed lines) interactions of HMGN2 molecule A (light purple) and HMGN2 molecule B (dark purple) with the H2A-H2B heterodimers on opposite faces of the nucleosome.

Both HMGN2 molecules use two major contact points in R23 and R26 at the ends of the acidic patch (Fig. 2; Structural basis of nucleosome recognition by the conserved Dsup and HMGN nucleosome-binding motif Supplemental Fig. S6). The binding of HMGN2 to the acidic patch is consistent with a proposed model for the interaction of HMGN2 with the nucleosome (Kato et al. 2011). Unexpectedly, however, our analysis revealed asymmetry between the two nucleosome-bound HMGN2 molecules. Specifically, while R23 in both HMGN2 molecules binds to the acidic patch through interactions with H2A E56 and H2B E113 (Fig. 2B; Supplemental Fig. S6), HMGN2 R26 interacts with H2A E61 and D90 in one HMGN2 molecule but with H2A E92 and D90 in the other HMGN2 molecule (Fig. 2B; Supplemental Fig. S6).

Previous work proposed that the binding of HMGN proteins to nucleosomes decreases via serine phosphorylation within the RRSARLSA region (Prymakowska-Bosak et al. 2001). To understand the effect of this modification upon HMGN2-nucleosome interactions, we modeled a phosphorylated version of HMGN2 S24 (pS24) in our cryo-EM maps; this modeling predicted an electrostatic repulsion with the acidic patch that would prevent the interactions seen with the unmodified protein (Supplemental Fig. S7). Additionally, we solved the cryo-EM structure of HMGN5 bound to a nucleosome. We observed that nucleosome binding by HMGN5, which is the most divergent HMGN family member (González-Romero et al. 2014), appears to be similar to that by HMGN2, and primarily involves the RRSAR motif of HMGN5 (Supplemental Fig. S8). It is also important to note that AI-based predictions with AlphaFold 3 (Abramson et al. 2024) generate only low confidence models of the HMGN2-bound nucleosome and do not identify the correct orientation of the region that binds to the acidic patch (Supplemental Fig. S9), as seen in our cryo-EM structure of HMGN2. Thus, HMGN-nucleosome complexes present a challenge for current AI-based structural prediction tools.

### The interactions of HMGN2 and Dsup with the acidic patch are related but not identical

The sequence and structural similarities between the nucleosome-binding motifs in Dsup and the HMGN proteins suggests a conserved mode of nucleosome recognition by these otherwise unrelated factors in *R. varieornatus* and humans, which have an estimated species divergence time of about 700 million years (Kumar et al. 2022). In this regard, Dsup R367 and HMGN2 R26 are arginine anchors that interact with the H2A triad E61, D90, and E92 (Supplemental Fig. S6; McGinty and Tan 2021), and their position within the nucleosome-binding domain align at the primary sequence level (Supplemental Fig. S6C). A distinction between the binding of HMGN2 and Dsup to the nucleosome is, however, the use of a slightly different register for the insertion of the type 1 arginine (Dsup R363 and HMGN2 R23) into the acidic patch via interactions with H2A E56 and H2B E113 (McGinty and Tan 2021). These findings suggest some degree of functional refinement of the nucleosome-binding domains of these factors during evolution.

### The 5S rDNA nucleosome exhibits conformations with closed (wrapped) and open (unwrapped) DNA ends

To gain a better understanding of the conformation and dynamics of nucleosomes containing a natural DNA sequence that is involved in gene activity, we conducted a cryo-EM analysis of the *5S rDNA* gene promoter nucleosome in the absence of HMGN proteins. We first obtained three structures from a non-crosslinked sample of the 167-bp 5S rDNA nucleosome, which contains a 10-bp linker DNA on each end (Fig. 3; Supplemental Figs. S10–S12 and Supplemental Table S1). The three 167-bp 5S rDNA nucleosome structures, which we refer to as closed, open I, and open II, lack density for the linker DNAs, and differ in the interactions involving the core nucleosomal DNA (Fig. 3; Supplemental Fig. S12). The closed 167-bp 5S rDNA nucleosome shows density for the entire 147-bp core DNA, which is wrapped around the histone octamer (Fig. 3A; Supplemental Fig. S12), and the open I and open II conformations show additional flexibility in one or both DNA ends, respectively (Fig. 3B,C; Supplemental Fig. S12). Thus, the 167-bp 5S rDNA nucleosome exhibits conformations with wrapped and unwrapped core DNA ends. To assess the effect of the 10-bp linkers on the Structural basis of nucleosome recognition by the conserved Dsup and HMGN nucleosome-binding motif flexibility of the core nucleosomal DNA in these structures, we next analyzed a sample of the 147-bp 5S rDNA nucleosome (Supplemental Fig. S13). In the absence of linkers, the 147-bp nucleosome core particle adopts the closed and open I conformations seen with the 167-bp 5S rDNA nucleosomes (Supplemental Fig. S14). We did not, however, observe a state analogous to open II, indicating that the flexibility of the core 5S rDNA increases in the presence of linker DNA. Notably, most nucleosomes in our 147-bp (∼60%) and 167-bp (∼95%) datasets adopt unwrapped conformations (Supplemental Figs. S10, S13), agreeing with the increased breathing dynamics of natural DNA sequences compared to the synthetic 601 DNA (Isaac et al. 2016).

**Figure 3.**
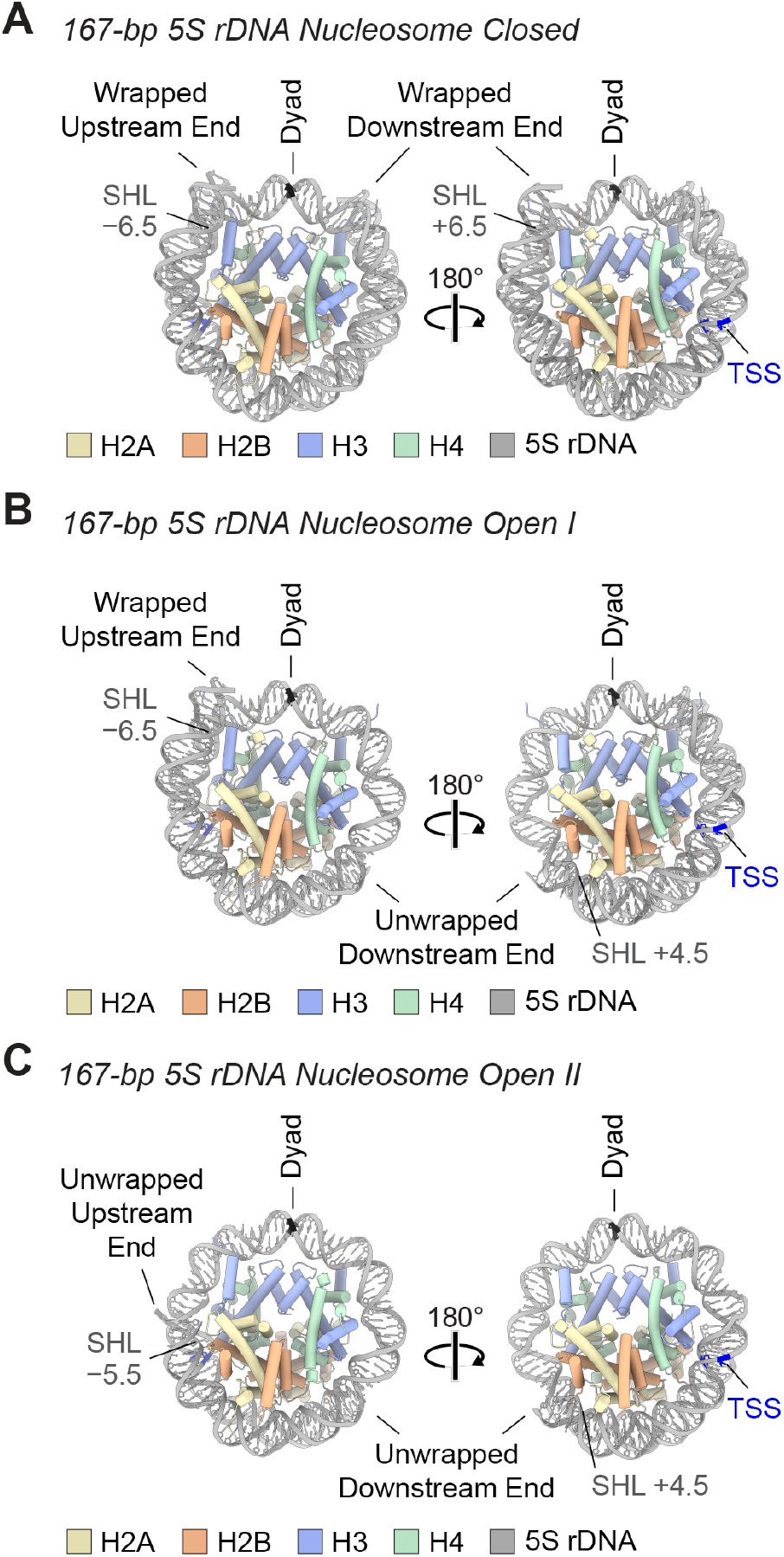
The 167-bp 5S rDNA nucleosome exhibits conformations with wrapped (closed) as well as unwrapped (open) DNA ends. Models of the 167-bp 5S rDNA nucleosome structures. (*A*) Closed conformation of the 167-bp 5S rDNA nucleosome (both DNA ends wrapped). (*B*) Open I conformation of the 167-bp 5S rDNA nucleosome (one wrapped end and one unwrapped end). (*C*) Open II conformation of the 167-bp 5S rDNA nucleosome (both DNA ends unwrapped). The 5S rDNA ends are labeled as upstream or downstream relative to the transcription start site (TSS). SHL: superhelical location.

### The DNA end that is downstream of the transcription start site (TSS) is preferentially unwrapped in the 5S rDNA nucleosome

Modeling of the 5S rDNA in the 147-bp and the 167-bp nucleosome structures revealed that DNA unwrapping is more prominent on the end that is downstream of the TSS (Fig. 3; Supplemental Figs. S11–S14). First, the downstream end lacks density in both the open I and open II 5S rDNA nucleosome conformations (Fig. 3B,C; Supplemental Figs. S12, S14), whereas the upstream end exhibits high flexibility only in the open II state (Fig. 3C; Supplemental Fig. S12). Second, the lack of DNA density is greater at the downstream end (beyond SHL +4.5 in open I and open II) relative to the upstream end (beyond SHL –6 in open II). Third, the preferential opening of the downstream DNA end is consistent with the higher number of particles observed in the open I classes compared to those detected in the other conformations of the 5S rDNA nucleosomes (Supplemental Figs. S10, S13). Finally, we did not detect nucleosomes that show flexibility exclusively at the upstream DNA end (Supplemental Figs. S10, S13). Increased flexibility in DNA that is downstream of the TSS may facilitate the transcription process, for instance, by exposing DNA binding sites for transcription factors (White 1998).

In addition to asymmetry in the flexibility of the DNA ends, we observed two arginines with different side chain conformations in the two H2A-H2B dimers of the 5S rDNA nucleosomes (Supplemental Figs. S15, S16). Specifically, R77 of H2A near the DNA end upstream of the TSS inserts into the minor groove at SHL –5.5, contributing to the stabilization of the DNA around the histone octamer in the closed, open I, and open II conformations. In contrast, in the two open states of the 5S rDNA nucleosomes, R77 of H2A near the downstream DNA end adopts a conformation that is not compatible with a fully wrapped DNA turn (Supplemental Figs. S15, S16). Interestingly, the loss of the interaction of H2A R77 with the minor grove has been correlated with gain of DNA flexibility by molecular dynamics simulation and cryo-EM analysis of nucleosome core particles (Kniazeva et al. 2022; Ding et al. 2024). Furthermore, R86 of H2B facing the downstream outer gyre of the nucleosome establishes fewer contacts with the DNA than R86 in H2B on the opposite side (Supplemental Fig. S15). Thus, we observe asymmetries in H2A R77 and H2B R86 on the opposite faces of the 5S rDNA nucleosome that mirror the different flexibility of the DNA ends.

### Visualization of 5S rDNA nucleosome dynamics

To visualize the dynamics of the 5S rDNA nucleosome, we performed principal component analysis (PCA) of the 167-bp nucleosome data set and then generated 3D movies with the frames corresponding to the first two components of conformational variability (van Heel and Frank, 1981; Punjani and Fleet, 2021). PCA revealed that the main source of Structural basis of nucleosome recognition by the conserved Dsup and HMGN nucleosome-binding motif variability corresponds to the attachment/detachment of the downstream 5S rDNA end to/from the histone octamer. The transition between the closed conformation and the unwrapping of the downstream DNA end is indicated by the fading of the density between SHL +5 and SHL +7 and the increased mobility of H3αN and H2A residues 107-116 in some frames (Supplemental Movie S1). Consistent with this analysis, correlations between stabilization/flexibilization of these histone regions and DNA wrapping/unwrapping have been suggested for nucleosomes reconstituted in vitro and nucleosomes isolated from cells (Luger et al. 1997; Ferreira et al. 2007; Shukla et al. 2011; Arimura et al. 2021). The main component in the PCA also captures a bulge in the upstream DNA end and histone flexibility on the opposite side of the nucleosome (Supplemental Movie S1). The second component of variability in the data mostly involves the flexibility of the DNA upstream of the TSS, as indicated by a fully wrapped upstream DNA that progressively unwraps from SHL −7 to SHL −6, while the flexible downstream DNA, which lacks density between SHL +5 and SHL +7, partially re-wraps around the histone octamer (Supplemental Movie S2). Altogether, the PCA is consistent with the preferential opening of the downstream DNA observed in the 5S rDNA nucleosome open I and open II structures.

### Crosslinking of the 5S rDNA nucleosomes results in DNA unwrapping

Glutaraldehyde (5 Å spacer arm) and formaldehyde (2-3 Å spacer arm) are often used to stabilize biomolecular interactions in cryo-EM samples containing nucleosomes (see, for example: Song et al. 2014; Wang et al. 2021; Lewis et al. 2021; Shioi et al. 2024). We set out to test their effects on the 5S rDNA nucleosome dynamics. Unexpectedly, crosslinking of the 147-bp and 167-bp nucleosomes with either glutaraldehyde or formaldehyde resulted only in cryo-EM structures with both ends unwrapped (Supplemental Figs. S17–S20), which is in contrast to non-crosslinked samples that show both closed and open conformations. Furthermore, the crosslinked nucleosomes exhibit DNA that is flexible starting at SHLs ±5 (Supplemental Fig. S20), indicating greater flexibility in the upstream DNA end in crosslinked nucleosomes.

We did not detect closed, open I, or open II states in either the 147-bp or the 167-bp crosslinked nucleosomes (Supplemental Figs. S17–S20), which suggests that crosslinking affects the inherent dynamics of 5S rDNA nucleosomes. Moreover, the increased flexibility in the upstream DNA end in the crosslinked relative to non-crosslinked nucleosomes correlates with a configuration of the side chain of H2A R77 that would collide with a fully wrapped DNA turn (Supplemental Fig. S21A). In addition, the two H2A R77 residues in the glutaraldehyde-crosslinked 167-bp nucleosome adopt different conformations (Supplemental Fig. S21B) that are consistent with the highly flexible DNA ends. These findings support a relationship between the conformation of H2A R77 and the flexibility of the nearby nucleosomal outer gyre.

### Summary and perspectives

In this study, we used cryo-EM to visualize the interaction of the conserved Dsup and HMGN nucleosome-binding motif with the nucleosome. We first examined the tardigrade-specific Dsup protein, which we had previously found to be a nucleosome-binding protein with an HMGN-like motif (Chavez et al. 2019). This work revealed that the HMGN-like segment in Dsup binds to the nucleosome via the acidic patch (Fig. 1). We then performed cryo-EM with the human HMGN2 protein, which is a canonical HMGN protein that is found in all vertebrates (González-Romero et al. 2014), and saw that the HMGN nucleosome-binding domain interacts with the acidic patch in a manner similar to Dsup (Fig. 2). The combined protein sequence and structural data thus show that the HMGN nucleosome-binding domain is an ancient protein motif that is conserved from tardigrades to humans. It thus appears that the HMGN nucleosome-binding domain existed prior to the evolutionary split between protostomes and deuterostomes, and there may be an untapped realm of chromatin proteins with HMGN-like nucleosome-binding domains.

To understand the structure and dynamics of a nucleosome containing a natural DNA sequence that is involved in gene activation, we explored the use of the promoter region of the *X. borealis 5S rDNA* gene for our cryo-EM analyses of the HMGN2-nucleosome complex (Fig. 2) as well as of the nucleosome alone (Fig. 3). We observed conformational rearrangements that are consistent with nucleosome dynamics that facilitate transcription (Eltsov et al. 2018; Tan et al. 2023). The 601 sequence, which has an unusually high affinity for the histone octamer (Lowary and Widom 1998), can be readily reconstituted into nucleosomes, but it does not provide a good model for natural chromatin. Thus, for nucleosome structural studies, the 5S rDNA sequence is an appealing alternate to the 601 sequence. Our analysis, which showed that crosslinkers affect the conformations of the nucleosomal 5S rDNA, suggests that caution should be taken when drawing conclusions about DNA dynamics in samples involving crosslinked nucleosomes.

There remains much to be learnt in the future. For instance, due to the generally high flexibility of Dsup and HMGN2 bound to the nucleosome, we have yet to identify any other key contact points between these proteins and the nucleosome; yet, at the same time, the flexibility of Dsup and HMGN2 on the nucleosome also likely explains why these abundant nucleosome-binding proteins are compatible with enzymes, such as DNA and RNA polymerases, that traverse nucleosomes. It would be interesting and informative to explore the properties of the HMGN-like nucleosome-binding domain beyond the canonical HMGN proteins. We also hope that the 5S rDNA nucleosome will be useful for future studies of chromatin structure, function, and dynamics. Structural basis of nucleosome recognition by the conserved Dsup and HMGN nucleosome-binding motif

## Acknowledgements

We thank Torrey Rhyne, George Kassavetis, and Marta Sanz Murillo for critical reading of the manuscript. The HMGN5 protein was a kind gift from Dr. George A. Kassavetis. J.T.K. is the Amylin Chair in the Life Sciences. This work was supported by NIH grants R35 GM118060 to J.T.K. and R35 GM145296 to A.E.L. G.C-B. is an UCSD Chancellor’s Fellow and NEB Fellow. The authors acknowledge the facilities of the UC San Diego cryo-EM facility, part of the Goeddel Family Technology Sandbox, along with the scientific and technical assistance of facility staff members Dr. Mariusz Matyszewski and Dr. Inga Kuschnerus. We thank the Physics Computing Facility for their technical support. This paper was typeset with the bioRxiv word template by @Chrelli: www.github.com/chrelli/bioRxiv-word-template.

## Author contributions

J.T.K., G.C-B., J.A-L., and A.E.L. initially conceived the project. A.E.L. and J.T.K oversaw the overall execution of this work. G.C-B purified the proteins and prepared the samples for the cryo-EM analyses. J.A-L. optimized sample vitrification, collected cryo-EM data, and performed image processing and model building. J.A-L., G.C-B., J.T.K., and A.E.L. prepared the figures and wrote the manuscript.

Structural basis of nucleosome recognition by the conserved Dsup and HMGN nucleosome-binding motif

## Data availability

Models and maps have been deposited in the Protein Data Bank and Electron Microscopy Data Bank and will be released after acceptance of the paper for peer-reviewed publication.

## Materials and methods

### Cryo-EM sample preparation

The Dsup-601 nucleosome complex and the HMGN-5S rDNA nucleosome complexes were reconstituted essentially as described in Chavez et al.

(2019), except that the nucleosomes contained recombinant human histones. The core histones, which include the H2A and H2B amino acid residues that contact Dsup, are conserved between humans and *R. varieornatus* (Supplemental Fig. S22). Samples containing nucleosomes (in the absence of Dsup or HMGN proteins), Dsup-bound nucleosomes, or HMGN-bound nucleosomes were incubated with the crosslinking reagent [glutaraldehyde at 0.25% (v/v) final concentration for 5 min on ice, or formaldehyde at 1% (v/v) final concentration for 10 min at 22 °C; both reagents are in buffer HE: 25 mM Hepes-K^+^, pH 7.6, and 0.1 mM EDTA], and the reactions were stopped by the addition of 1 M Tris-HCl, pH 8.0, to 50 mM final concentration. In addition, non-crosslinked samples were prepared in parallel with buffer HE only in the reactions. The crosslinked and non-crosslinked samples were kept on ice prior to use in grid preparation for cryo-EM analysis. Additional methods are included in the Supplemental Material.

## SUPPLEMENTAL MATERIAL

**Supplementary Figure S1.**
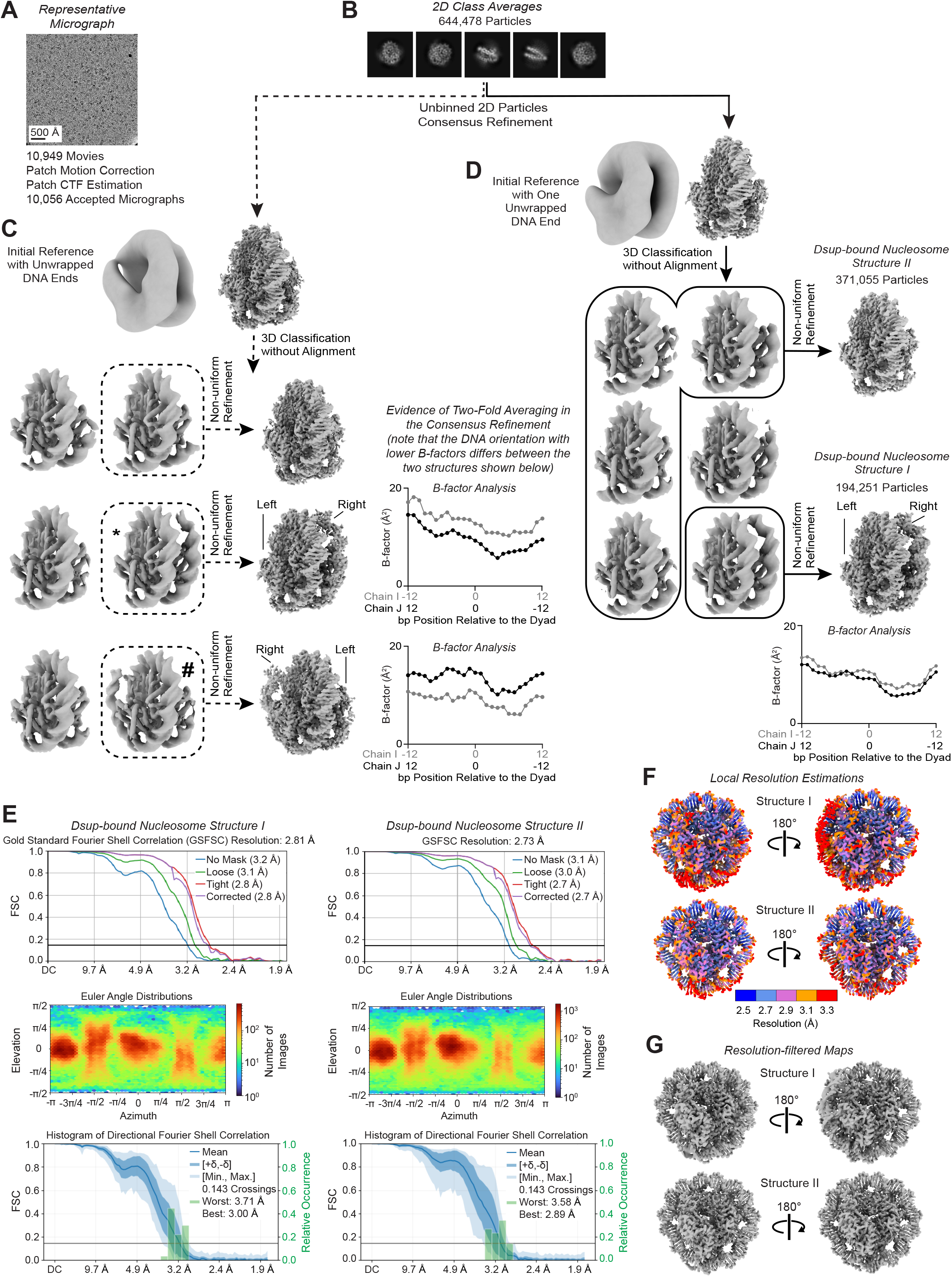
Image processing of the crosslinked Dsup-bound nucleosomes. Data collected from a single cryo-EM grid containing a crosslinked Dsup-nucleosome complex sample resulted in two structures, which we term structure I and structure II. (A) Representative micrograph, low-pass filtered to 10 Å. (B) Selected 2D class averages. (C) 3D refinement after 2D classifications using a low-pass filtered nucleosome with unwrapped DNA ends as the initial reference. Subsequent 3D classification without alignment shows classes with apparent differential flexibility in opposite DNA ends (indicated with * and #). Per-bp B-factor analysis of the two possible DNA orientations (lower B-factor orientation inverted between plots). (D) 3D refinement using a low-pass filtered nucleosome with one DNA end wrapped around the octamer as the initial reference. Subsequent 3D classification without alignment shows preferential opening of the Dsup-bound nucleosome on the ‘left’ side, more precisely describing the dynamics in our dataset. (E) Fourier shell correlations (FSCs), particle angle distributions, and histograms of directional FSC. (F) Local resolution estimation in Å plotted in five different colors onto the refined maps. (G) Maps filtered according to the local resolution estimations.

**Supplementary Figure S2.**
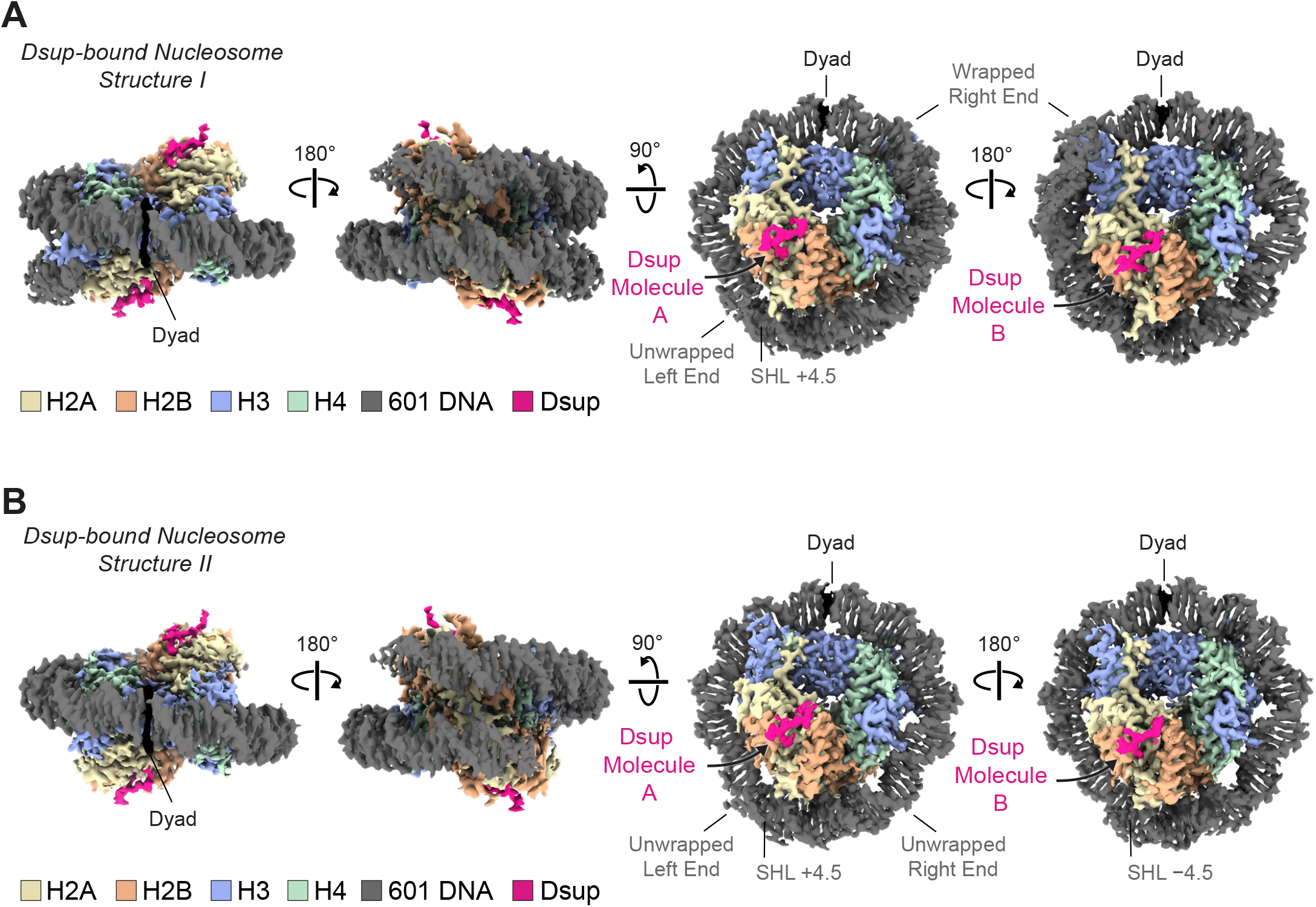
Cryo-EM maps of the Dsup-bound nucleosome structures. (A) Dsup-bound nucleosome structure I. (B) Dsup-bound nucleosome structure II. Each of the two structures shows density for two nucleo-some-bound Dsup molecules (Dsup molecule A and Dsup molecule B). Structure I has one wrapped DNA end and one unwrapped DNA end. Structure II has both DNA ends unwrapped. We refer to the 5’ end and the 3’ end of the 601 DNA sequence in these structures as the “left” and “right” ends, respectively. SHL: superhelical location.

**Supplementary Figure S3.**
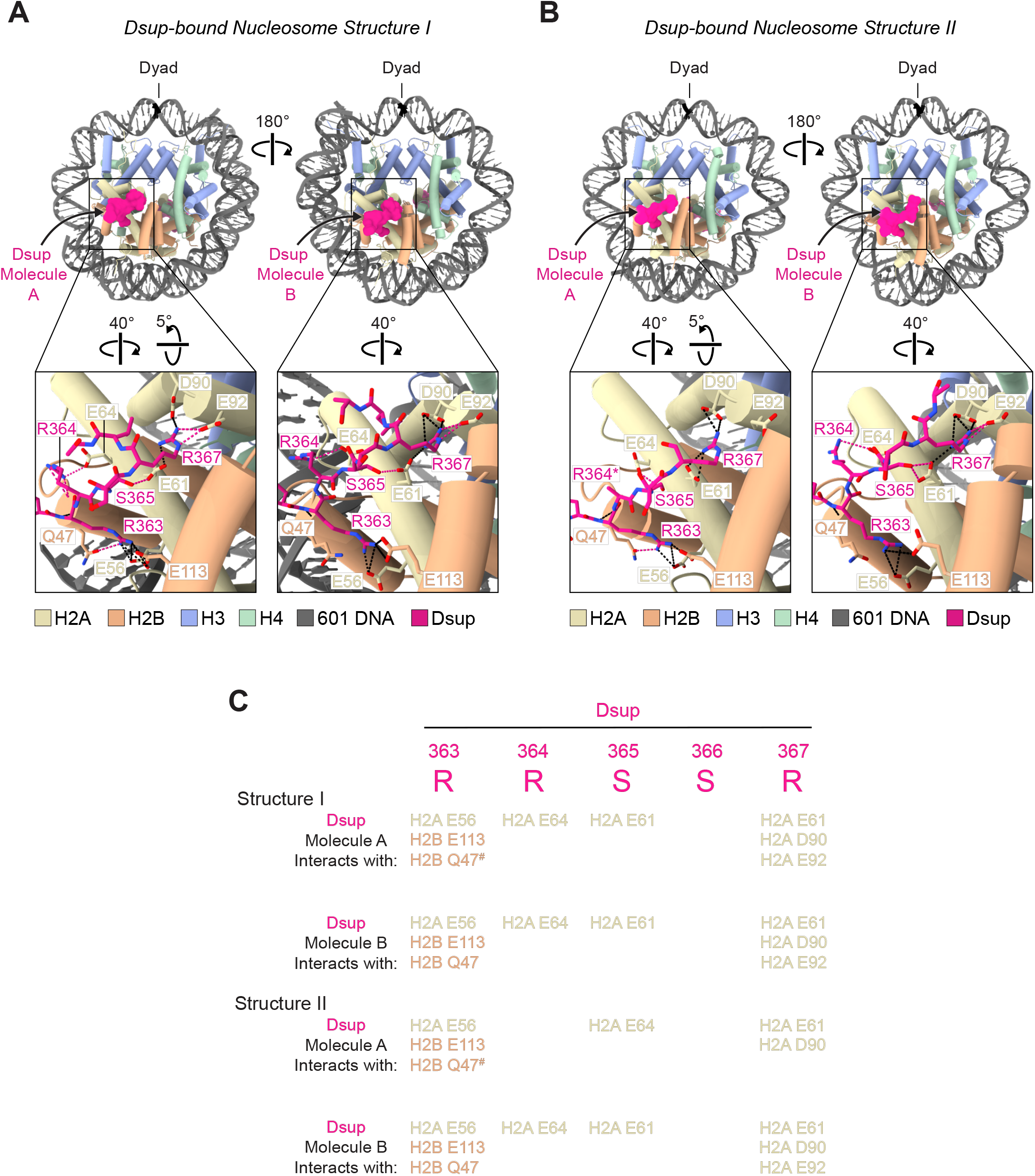
Interactions of Dsup with the nucleosome acidic patch and with Q47 of H2B. (A) Atomic model of the Dsup-bound nucleosome structure I. (B) Atomic model of the Dsup-bound nucleosome structure II. The black dashed lines in the inset images represent the interactions that are shared by Dsup molecule A and Dsup molecule B in structure I and structure II. The other Dsup interactions are depicted by pink dashed lines. The asterisk at R364 of one Dsup molecule indicates that the side chain is not modeled due to low resolution. (C) Summary of the interactions of Dsup molecule A and Dsup molecule B (residues 363-367) with the nucleosome acidic patch and with Q47 of the H2B α1-L1 elbow. The # symbol highlights the double hydrogen bonding between H2B Q47 and Dsup R363.

**Supplementary Figure S4.**
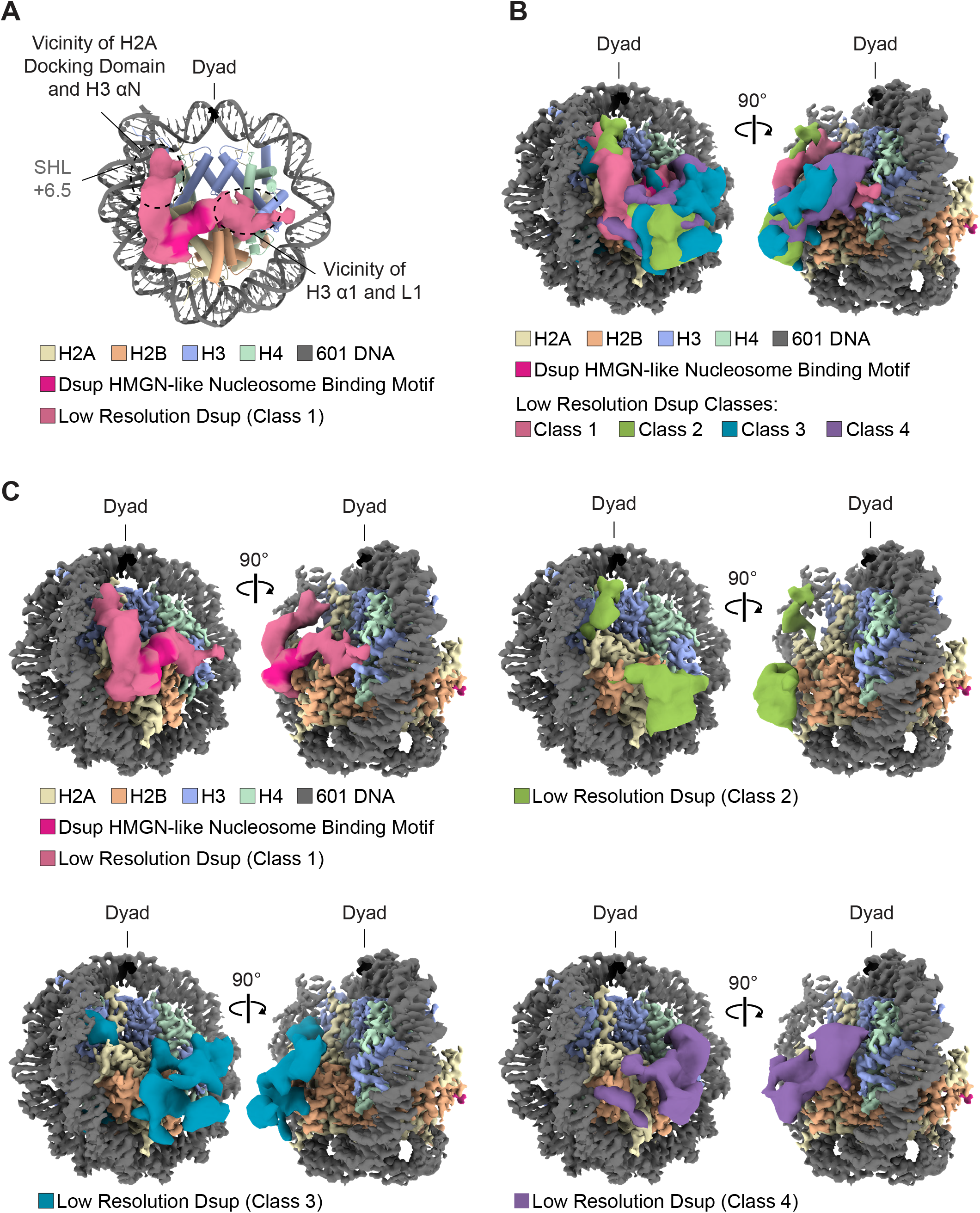
Dsup adopts different conformations beyond the nucleosome-binding core region. (A) Low-resolution Dsup density (class 1) obtained after nucleosome signal subtraction and focused classifications without alignment. SHL: superhelical location. (B) Overlay of four low-resolution electron microscopy Dsup classes. (C) Density maps of each of the four low-resolution Dsup classes.

**Supplementary Figure S5.**
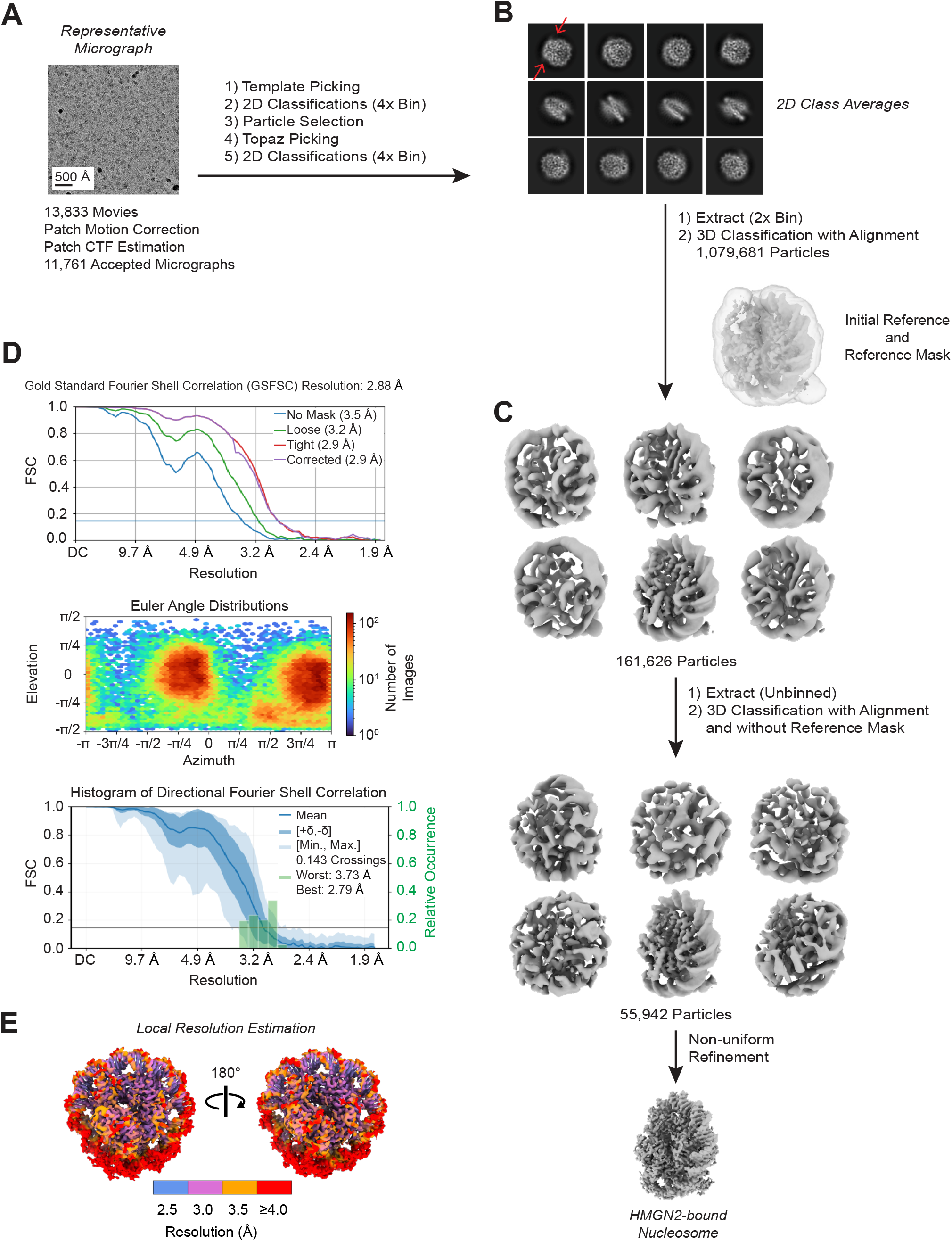
Image processing of the crosslinked HMGN2-bound nucleosomes. (A) Representative micrograph, low-pass filtered to 10 Å. (B) Selected 2D class averages. The two red arrows indicate the flexibility (absence of density) of both DNA ends. (C) Image processing scheme. (D) Fourier shell correlation (FSC), particle angle distribution, and histogram of directional FSC. (E) Local resolution estimation in Å plotted in four different colors onto the refined map.

**Supplementary Figure S6.**
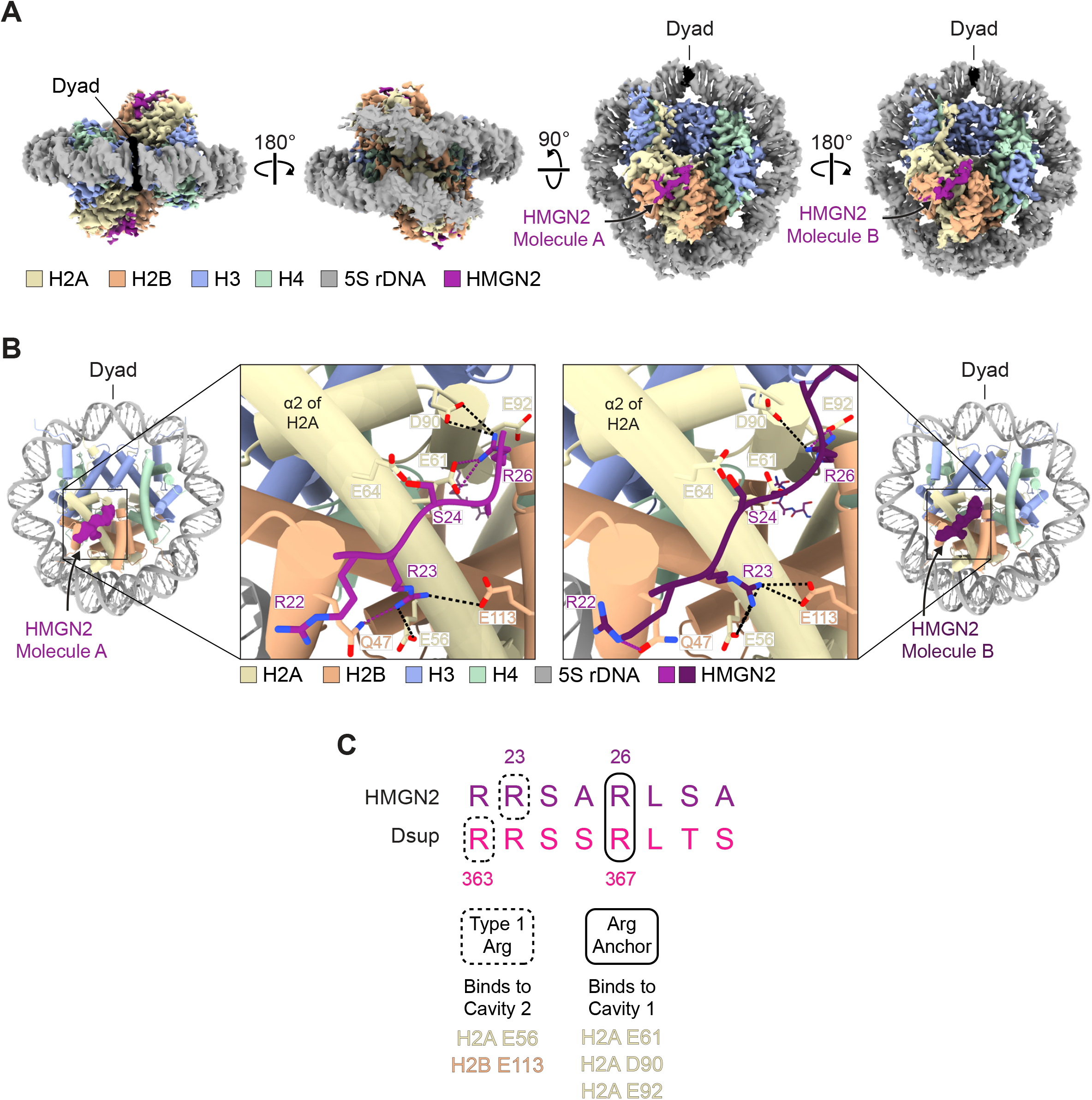
Interactions of HMGN2 with the nucleosome acidic patch and with Q47 of H2B. (A) Cryo-EM map of the HMGN2-bound nucleosome structure. (B) Model of the HMGN2-bound 167-bp 5S rDNA nucleosome. The close-ups highlight the interactions of HMGN2 molecule A and HMGN2 molecule B with the acidic patch and with H2B Q47. The black and purple dashed lines depict interactions that are shared and distinct, respectively, on each of the two faces of the nucleosome. (C) The interactions of HMGN2 and Dsup with the acidic patch are related but not identical. The HMGN2 and Dsup arginine anchor and Type 1 arginine show conserved and mismatched registers. Both the acidic patch and Q47 in the H2B α1-L1 elbow are hotspots for tethering nucleo-some-binding factors via interactions with arginine residues (McGinty and Tan 2021).

**Supplementary Figure S7.**
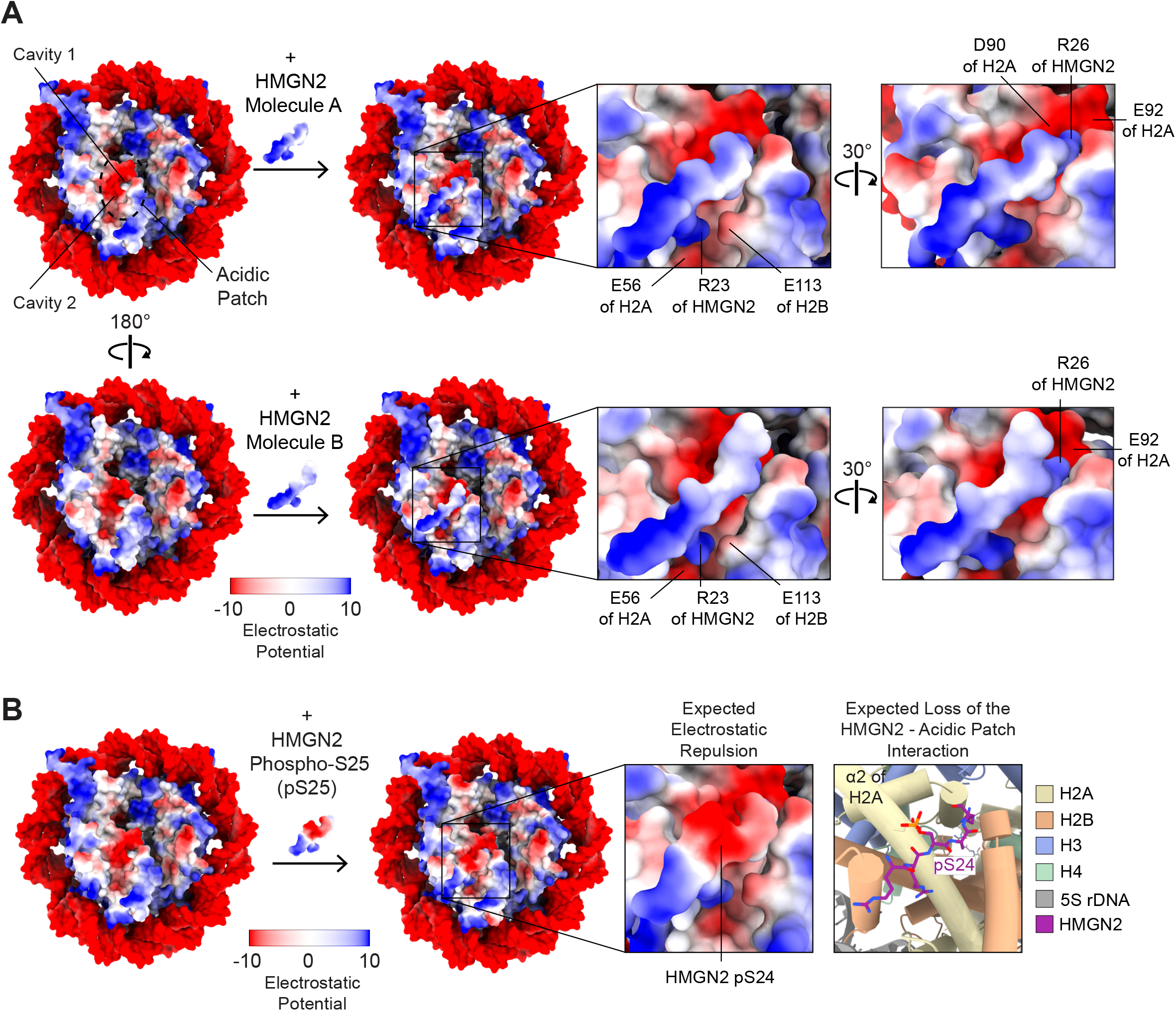
Charge distribution maps of the HMGN2-bound nucleosome. (A) HMGN2-bound 5S rDNA nucleosome colored by Coulombic electrostatic potential. In the farthest left surface images, the densities for HMGN2 molecule A and HMGN2 molecule B are removed from the maps for clarity. (B) The RRSAR segment of HMGN2 was modeled with phosphorylated S24 (pS24) to show the anticipated electrostatic repulsion with the negatively charged acidic patch (compare panels A and B).

**Supplementary Figure S8.**
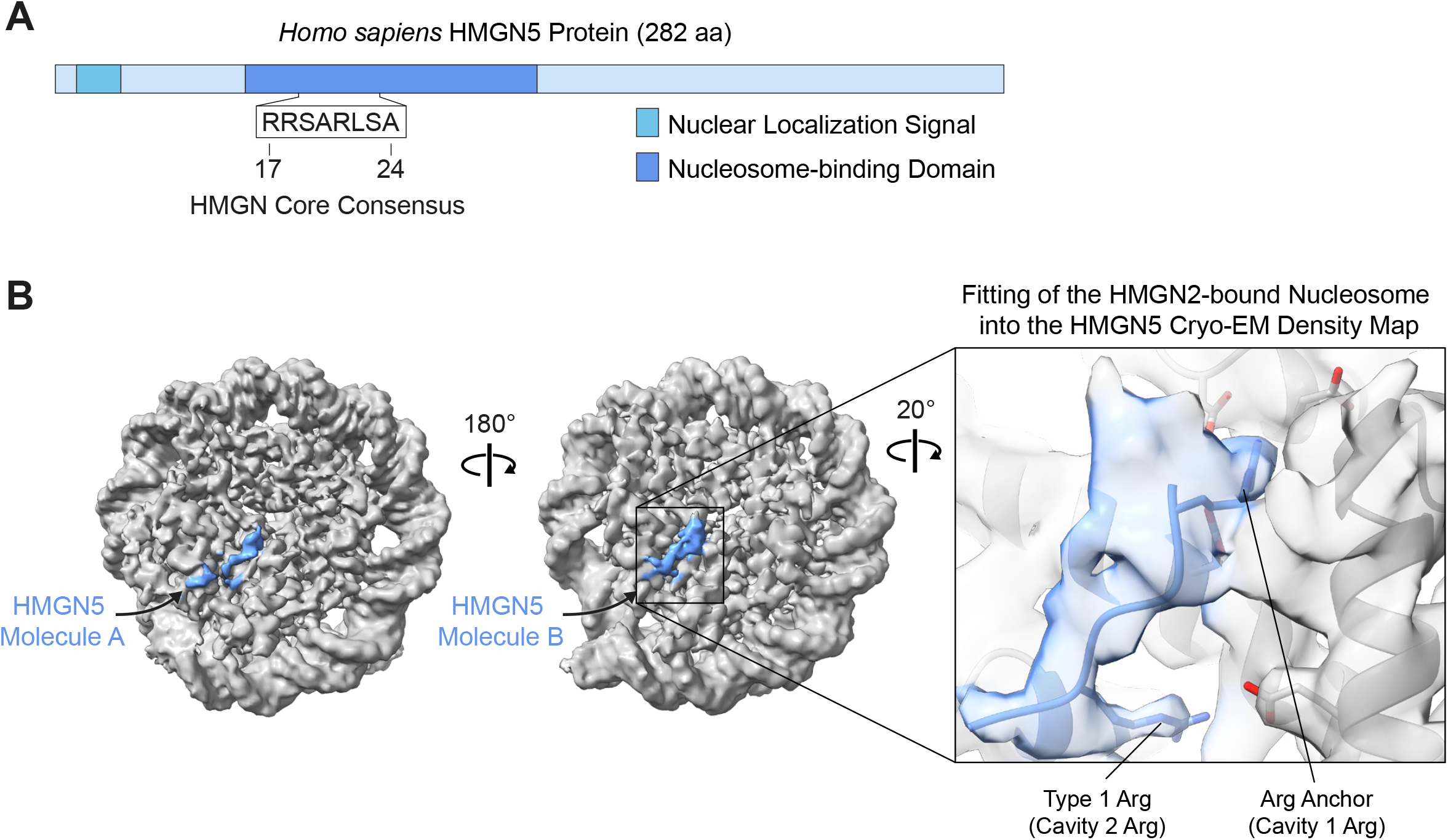
Binding to the nucleosome by HMGN5 appears to be similar to that by HMGN2. (A) H. sapiens HMGN5 protein. The sequence corresponds to the core consensus motif that anchors the HMGN proteins to the nucleosome (HMGN5 residues 17-24). (B) Cryo-EM map of the HMGN5-bound 167-bp 5S rDNA nucleosome crosslinked with glutaraldehyde. The close-up view on the right shows the model of the nucleosome bound to HMGN2 fitted into the HMGN5-nucleosome map.

**Supplementary Figure S9.**
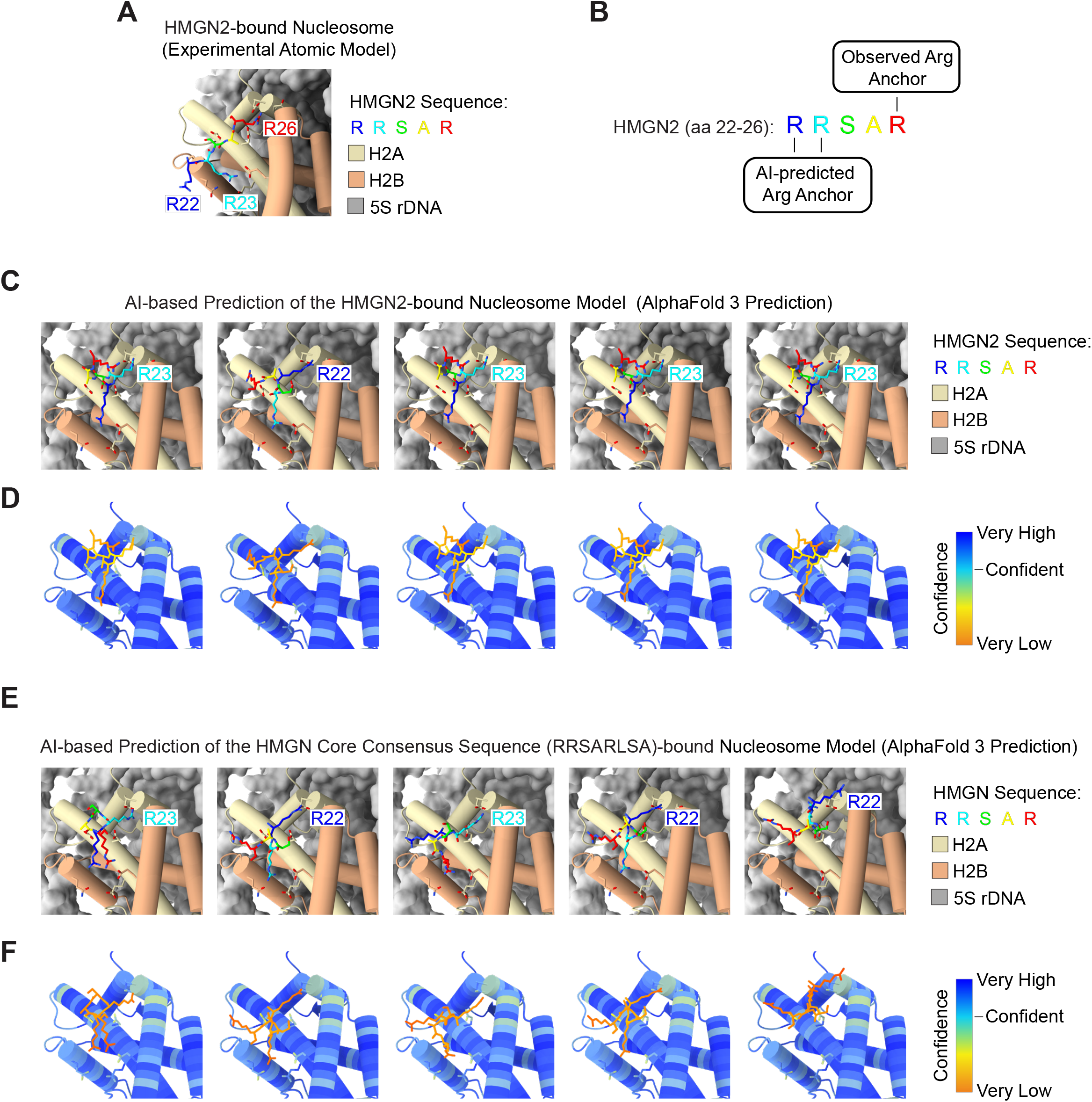
AI-based structural predictions do not correctly identify the experimentally observed arginine (Arg) anchor in HMGN2. (A) Experimentally determined structural model of HMGN2 bound to the nucleosome acidic patch. (B) Cryo-EM-observed and AI-predicted Arg anchor in HMGN2. (C,D) AI predictions of nucleosome-bound full-length HMGN2 with AlphaFold 3 (Abramson et al. 2024). (E,F) AI predictions of the nucleo-some-bound HMGN core consensus sequence (RRSARLSA) with AlphaFold 3. In panels A, C, and E, only the RRSAR sequence [rainbow-colored from N-to C-terminus (blue to red)] is shown in the models. Panels D and F display the per-residue accuracy of the models shown in C and E, indicated by the predicted Local Distance Difference Test (pLDDT): very low (< 50, orange), low (50-70, yellow), confident (70-90, cyan) and very high (> 90, blue). The predictions are mostly very high confidence for the histones and very low confidence for the HMGN motif.

**Supplementary Figure S10.**
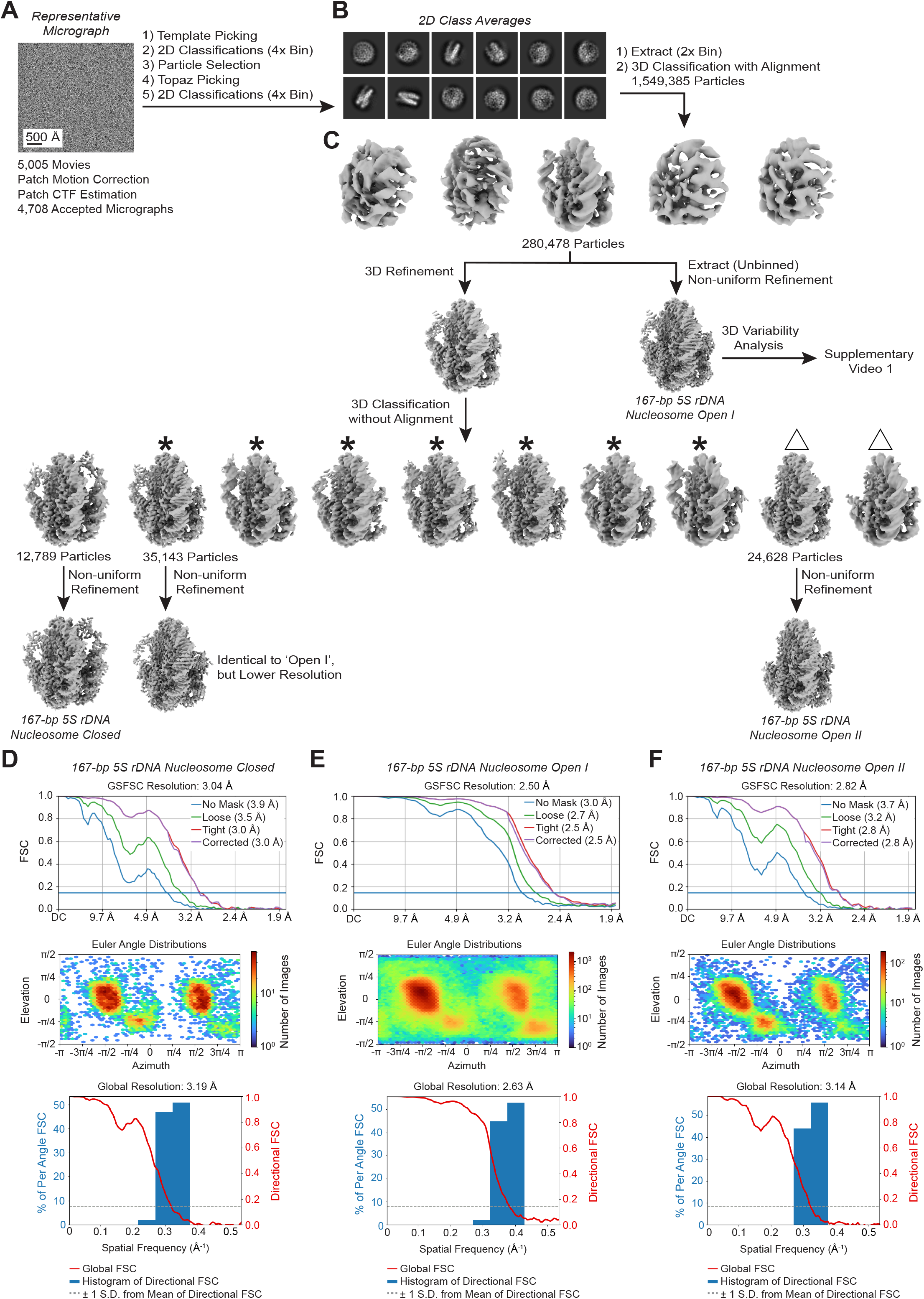
Image processing of the 167-bp 5S rDNA nucleosomes. (A) Representative micrograph, 10 Å low-pass filtered. (B) Selected 2D class averages. (C) 3D classifications and refinements. Classes marked with asterisks and triangles are 167-bp 5S rDNA nucleosome open I-like and open II-like, respectively. (D–F) Fourier shell correlations (FSC), particle angle distributions, and histogram of directional FSC for the 167-bp 5S rDNA nucleosome maps in the closed, open I, and open II conformations.

**Supplementary Figure S11.**
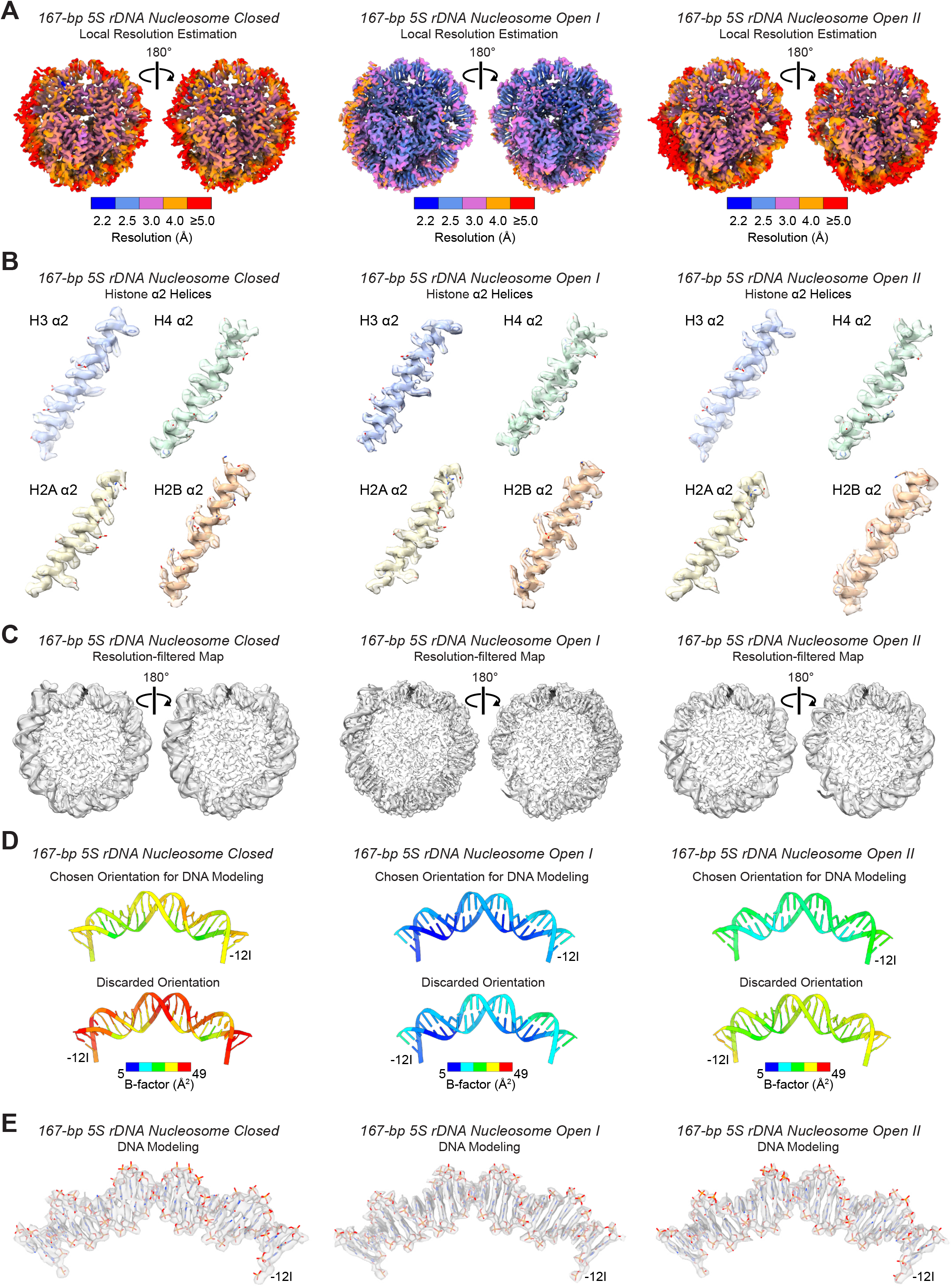
Local resolution estimation, map features, and DNA orientation analysis of the 167-bp 5S rDNA nucleosome structures. (A) Local resolution estimations plotted onto the maps. (B) Cryo-EM densities and models of the α2 helices of the histones in the three 167-bp 5S rDNA nucleosome states. (C) Maps filtered according to the local resolution estimation. The modeled DNA is shown as a ribbon. Histone models are removed from the map for clarity. We built a model for the DNA whenever density was observed in a map that had been filtered according to the estimated local resolution, without lowering the default display contour levels in ChimeraX (Goddard et al. 2018). (D) DNA orientation analysis. To distinguish between the two possible orientations of the modeled DNA (related through a 180° rotation along the dyad axis), we refined the 25 bp of nucleosomal DNA centered around the dyad (position 0) in those two orientations. The 25 bp are colored by the per-bp B-factors (see Supplemental Materials and methods). Lower B-factors indicate higher certainty about the atomic 3D coordinates. For the three structures, the B-factors are consistently lower for the same orientation. (E) Cryo-EM densities and chosen models of the 25-bp DNA centered around the dyad. We chose this DNA segment because of its high-resolution features in the three 167-bp 5S rDNA nucleosome maps.

**Supplementary Figure S12.**
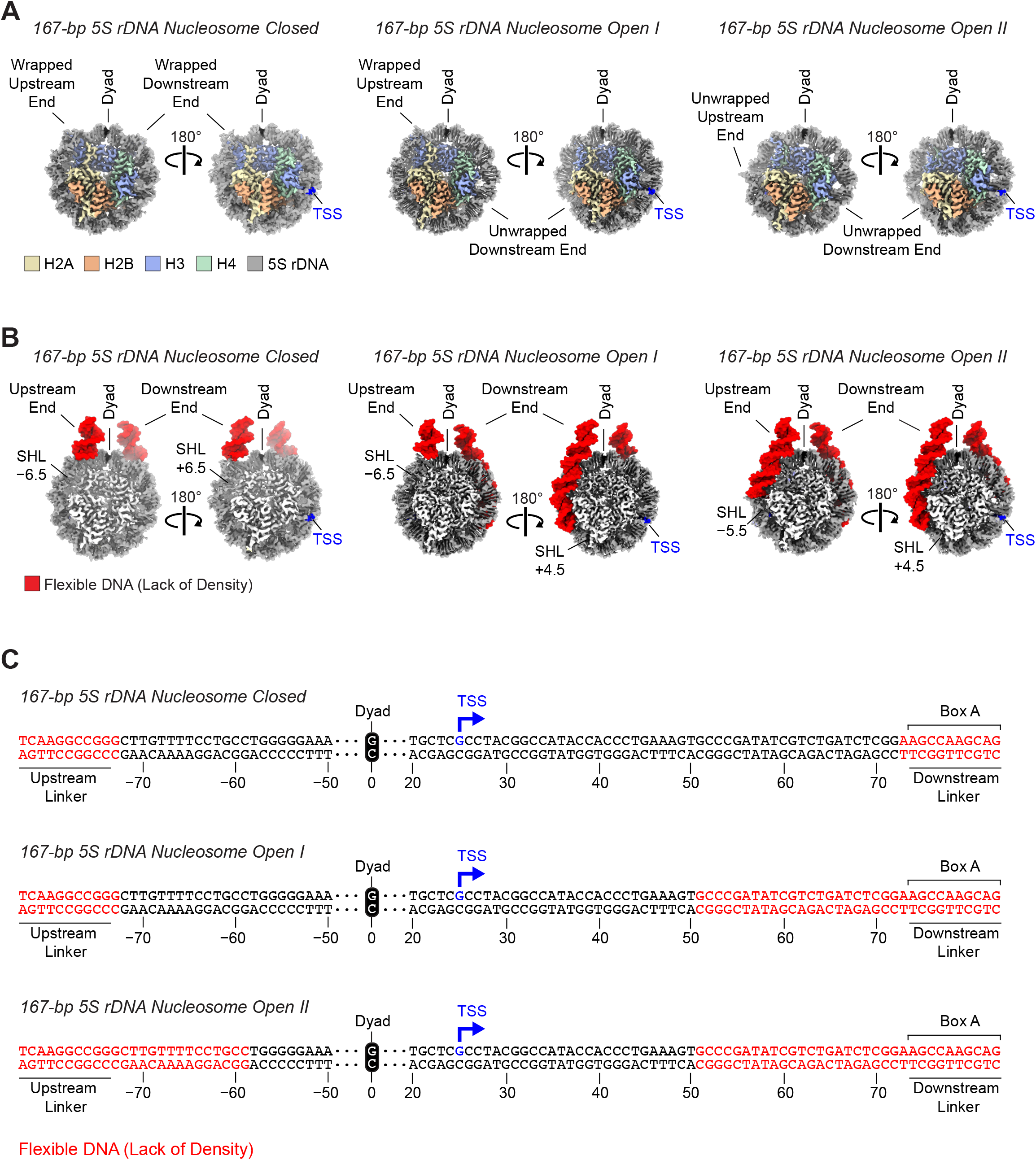
The DNA end that is downstream of the 5S rDNA TSS is preferentially unwrapped in the 167-bp nucleosome. (A) Cryo-EM maps of the 167-bp 5S rDNA nucleosome in the closed, open I, and open II conformations. (B) Cryo-EM maps with the addition of molecular models, in red, to indicate the DNA segments that lack cryo-EM density (flexible DNA). SHL: superhelical location. (C) Flexible DNA sequence in the 167-bp 5S rDNA nucleosomes. The DNA sequence in red type is not visible in the cryo-EM maps. The downstream linker sequence (AGCCAAGCAG) corresponds to most of the Box A sequence (AGCCAAGCAGGG) in the promoter of 5S rRNA genes (Pieler et al. 1987). The numbers indicate the bp position relative to the dyad. The full 167-bp 5S rDNA sequence is available in the Supplemental Materials and methods. TSS: transcription start site.

**Supplementary Figure S13.**
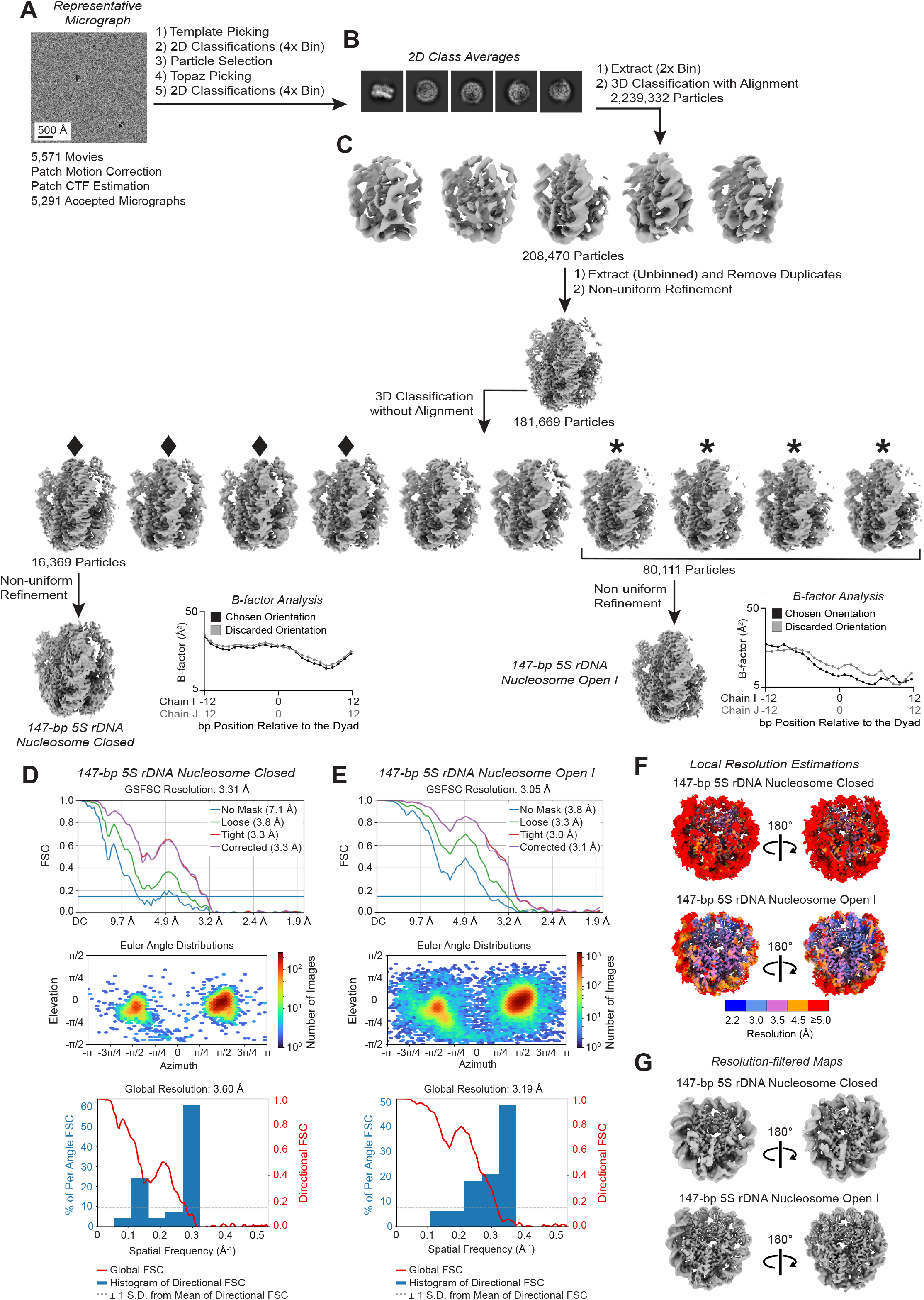
Image processing of the 147-bp 5S rDNA nucleosomes. (A) Representative aligned micrograph, 10 Å low-pass filtered. (B) Selected 2D class averages. (C) 3D classifications and refinements. Classes marked with diamonds and asterisks are 147-bp 5S rDNA nucleosome closed-like and open I-like, respectively. (D–E) Fourier shell correlations (FSC), particle angle distributions, and histogram of directional FSC for the 147-bp 5S rDNA nucleosome maps. (F) Local resolution estimations plotted onto the maps. (G) Maps filtered according to the local resolution estimation.

**Supplementary Figure S14.**
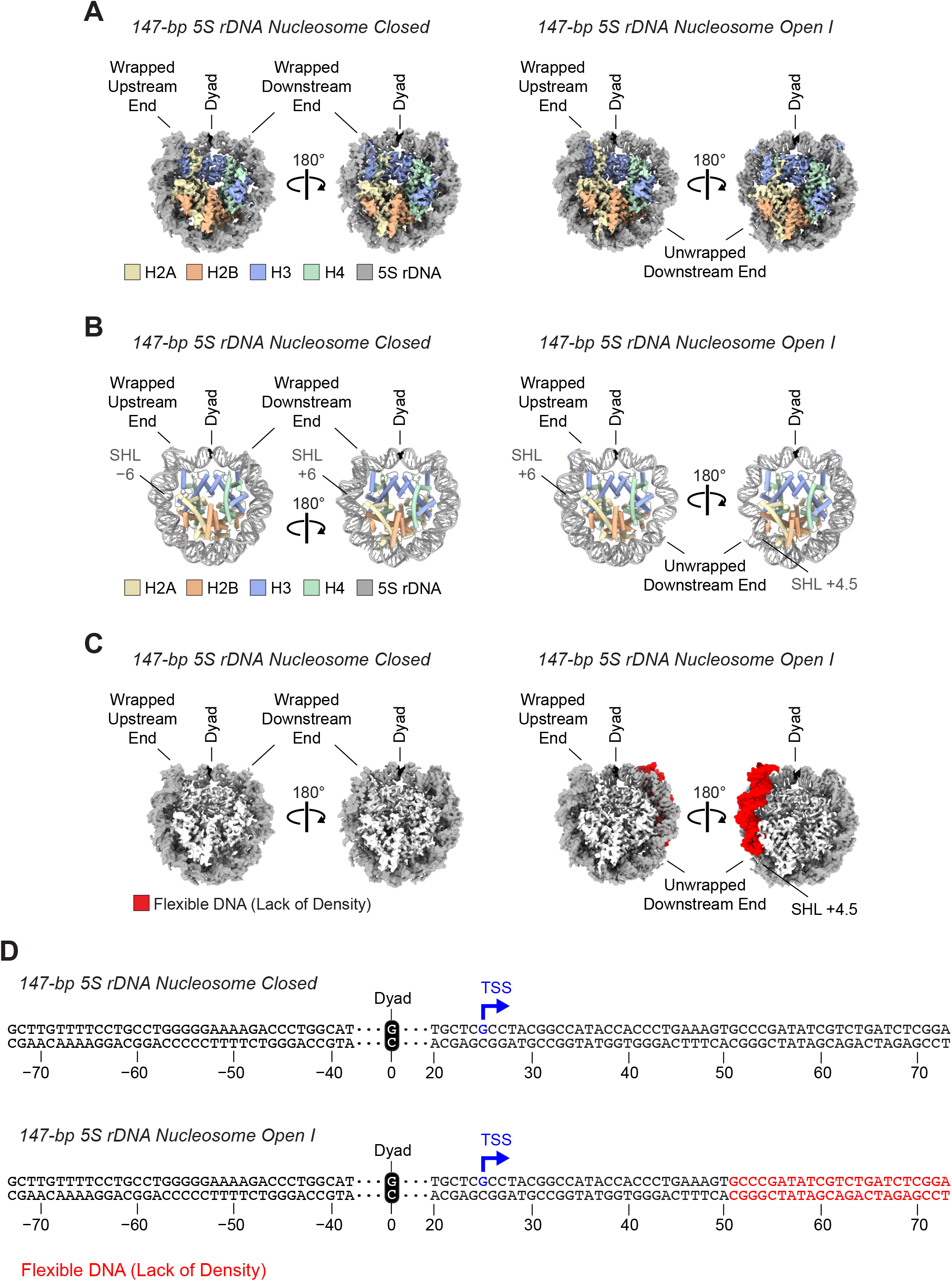
The 147-bp 5S rDNA nucleosome exists in a closed form as well as in an open form that is unwrapped downstream of the TSS. (A) Cryo-EM maps of the 147-bp 5S rDNA nucleosome in the closed (both DNA ends wrapped) and open I (one wrapped and one unwrapped DNA end) conformations. (B) Models built on the cryo-EM maps of the 147-bp 5S rDNA nucleosome structures. SHL: superhelical location. (C) Cryo-EM maps with the addition of red molecular models, which indicate the DNA segments that lack cryo-EM density. (D) Flexible DNA sequence in the 147-bp 5S rDNA nucleosome. The DNA sequence in red type is not visible in the cryo-EM map of the 147-bp 5S rDNA nucleosome open I conformation. The numbers indicate the bp position relative to the dyad. The 5S rDNA ends in panels A–C are labeled as upstream or downstream relative to the transcription start site (TSS). The full 147-bp 5S rDNA sequence is available in the Supplemental Materials and methods.

**Supplementary Figure S15.**
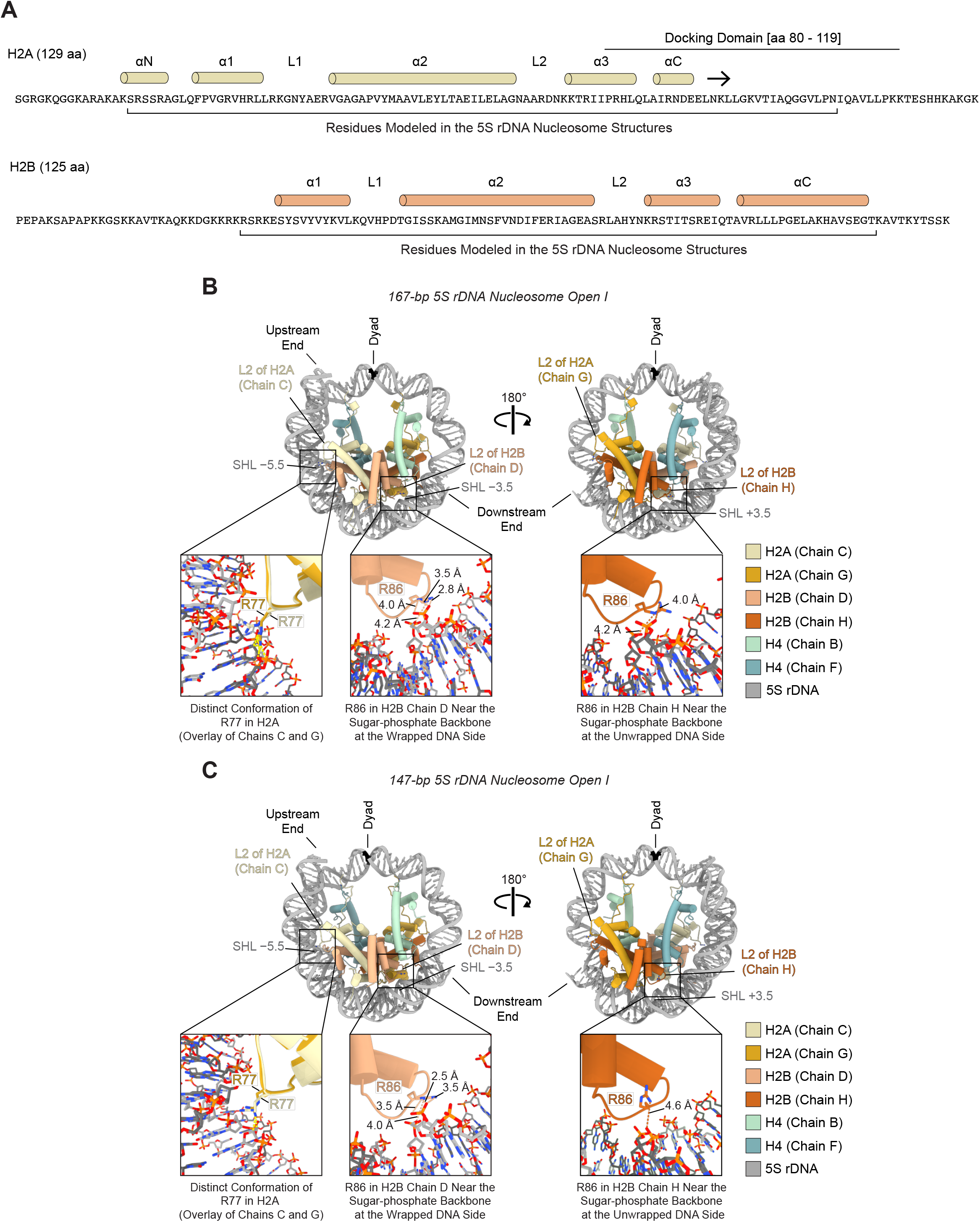
Distinct conformations of H2A R77 and H2B R86 in their interactions with wrapped versus unwrapped 5S rDNA in the nucleosome. (A) Sequence and secondary structure of H2A and H2B. α-helices and β-strands are represented by cylinders and arrows. L: Loop; N: N-terminal; C: C-terminal. The brackets highlight the histone sequence that is modeled in the structures of the 167-bp and the 147-bp 5S rDNA nucleosomes. (B,C) Models of the 167-bp and the 147-bp 5S rDNA nucleosomes in the open I conformation (one wrapped DNA end and one unwrapped DNA end) with close-up views of H2A R77 and H2B R86 on the two faces of the nucleosomes. Histone H3 is removed from the models for clarity. In the close-up images on the left, H2A (chain C) near the upstream DNA end is overlaid onto H2A (chain G) near the downstream DNA end to show that H2A R77 facing the flexible downstream end would clash (yellow dashed lines) with a fully wrapped outer DNA turn. The close-up views of H2B R86 on each of the two nucleosome faces (middle and right inset images) depict anticipated electrostatic interactions with the sugar-phosphate backbone of the DNA. The numbers indicate the distance (in Å) between H2B R86 and the sugar phosphate backbone. SHL: superhelical location.

**Supplementary Figure S16.**
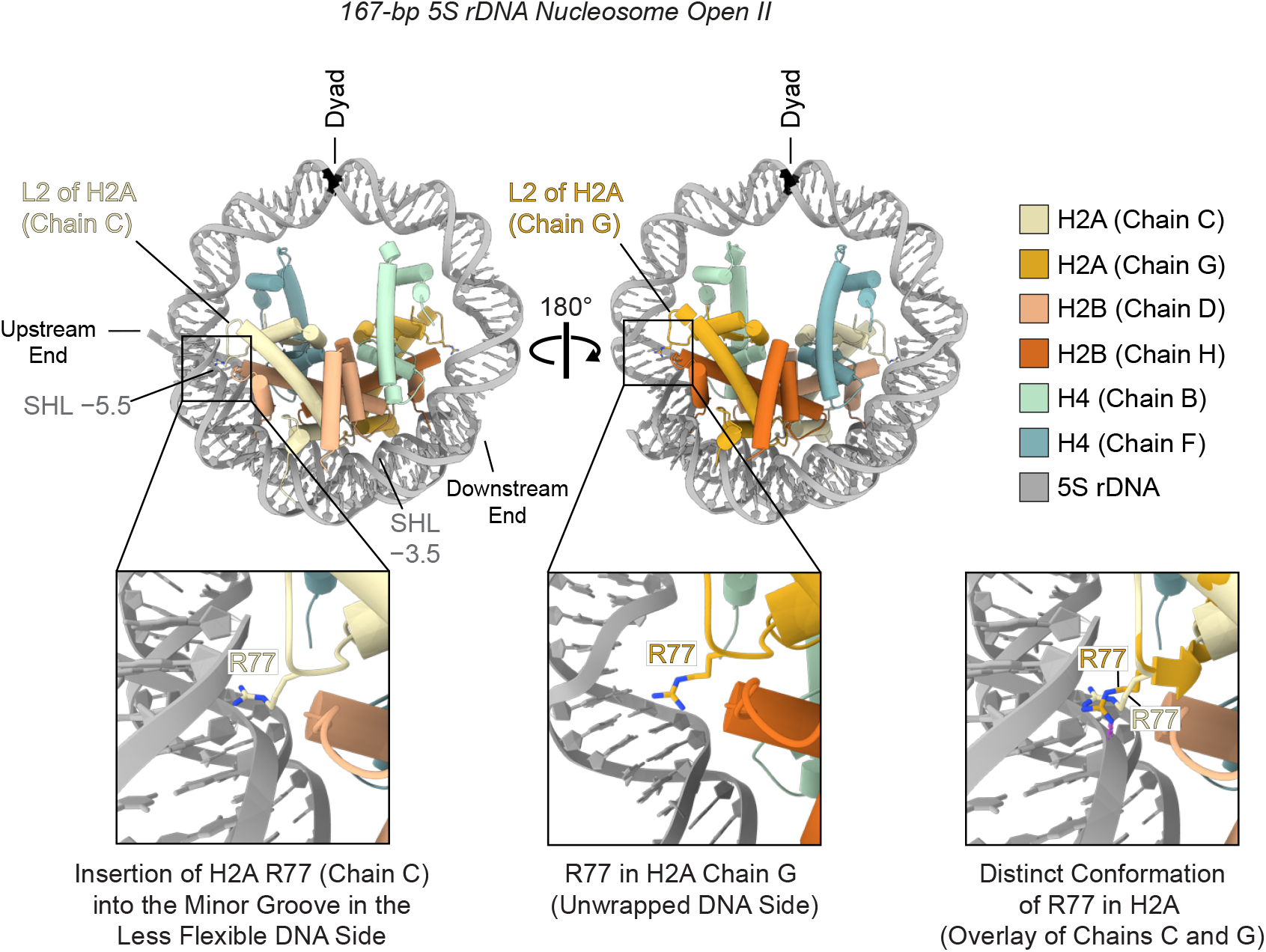
H2A R77 inserts into the minor groove in one side of the 167-bp 5S rDNA nucleosome in the open II conformation. Model with close-up views of H2A R77 on each of the two nucleosome faces. Histone H3 is removed from the model for clarity. R77 of H2A chain C interacts with the 5S rDNA near to the SHL −5.5 at the less flexible DNA side. In contrast, R77 of H2A chain G (near the most flexible downstream DNA side) would collide (depicted by purple dashed lines in the inset image on the right) with DNA wrapped at the SHL +5.5. SHL: superhelical location. Alegrio-Louro et al. (Leschziner), Supplemental Fig. S16

**Supplementary Figure S17.**
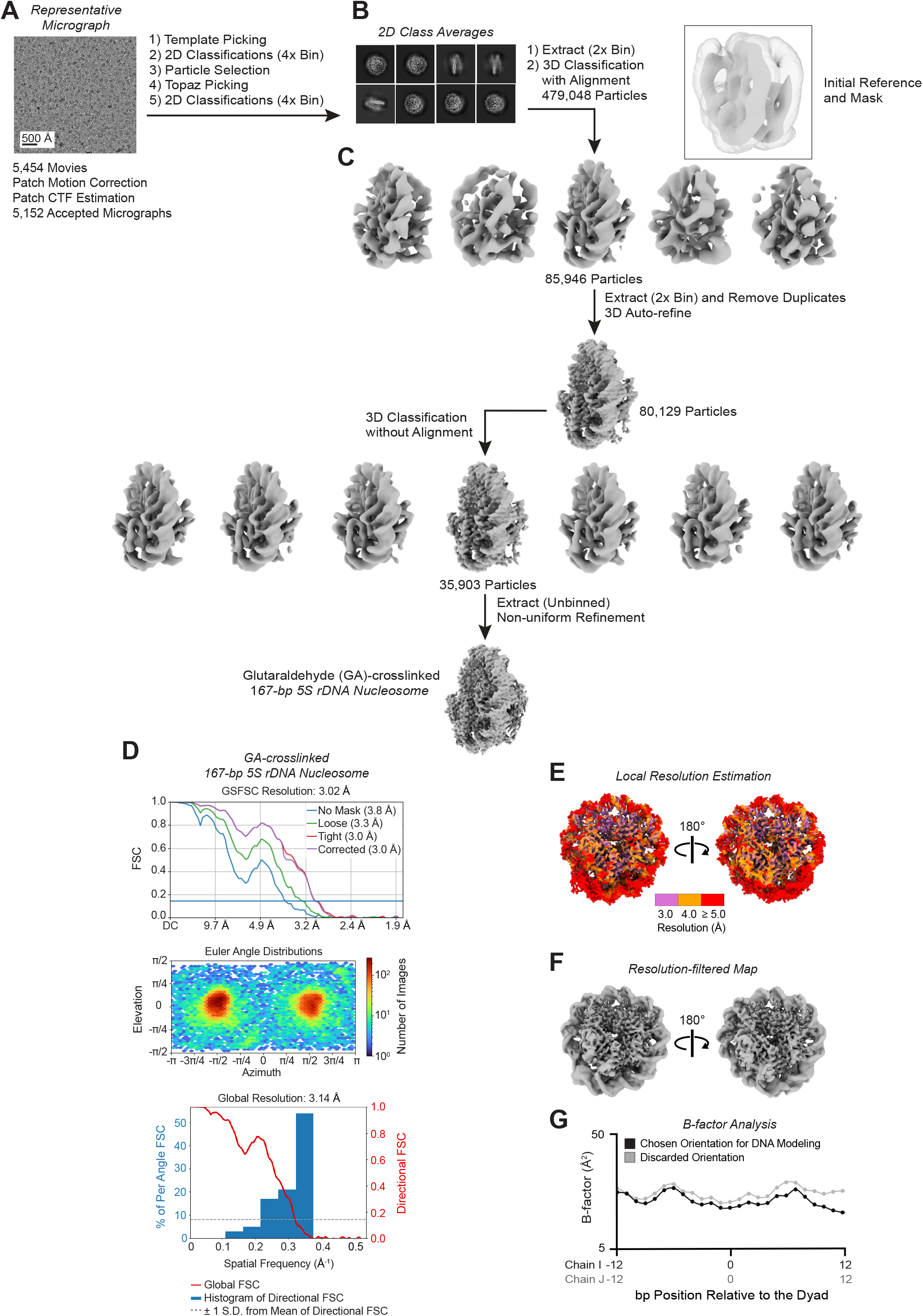
Image processing of the glutaraldehyde-crosslinked 167-bp 5S rDNA nucleosomes. (A) Representative micrograph, 10 Å low-pass filtered. (B) Selected 2D class averages. (C) 3D classifications and refinements. Note that the initial reference for 3D classifications with alignment presented one DNA end wrapped around the octamer but the 3D classification output volume displayed both ends flexible. (D) Fourier shell correlation (FSC), particle angle distribution, and histogram of directional FSC. (E) Local resolution estimation plotted onto the map. (F) Map filtered according to the local resolution estimation. (G) Average per-bp B factors plotted for the 25 bp refined in the two possible orientations.

**Supplementary Figure S18.**
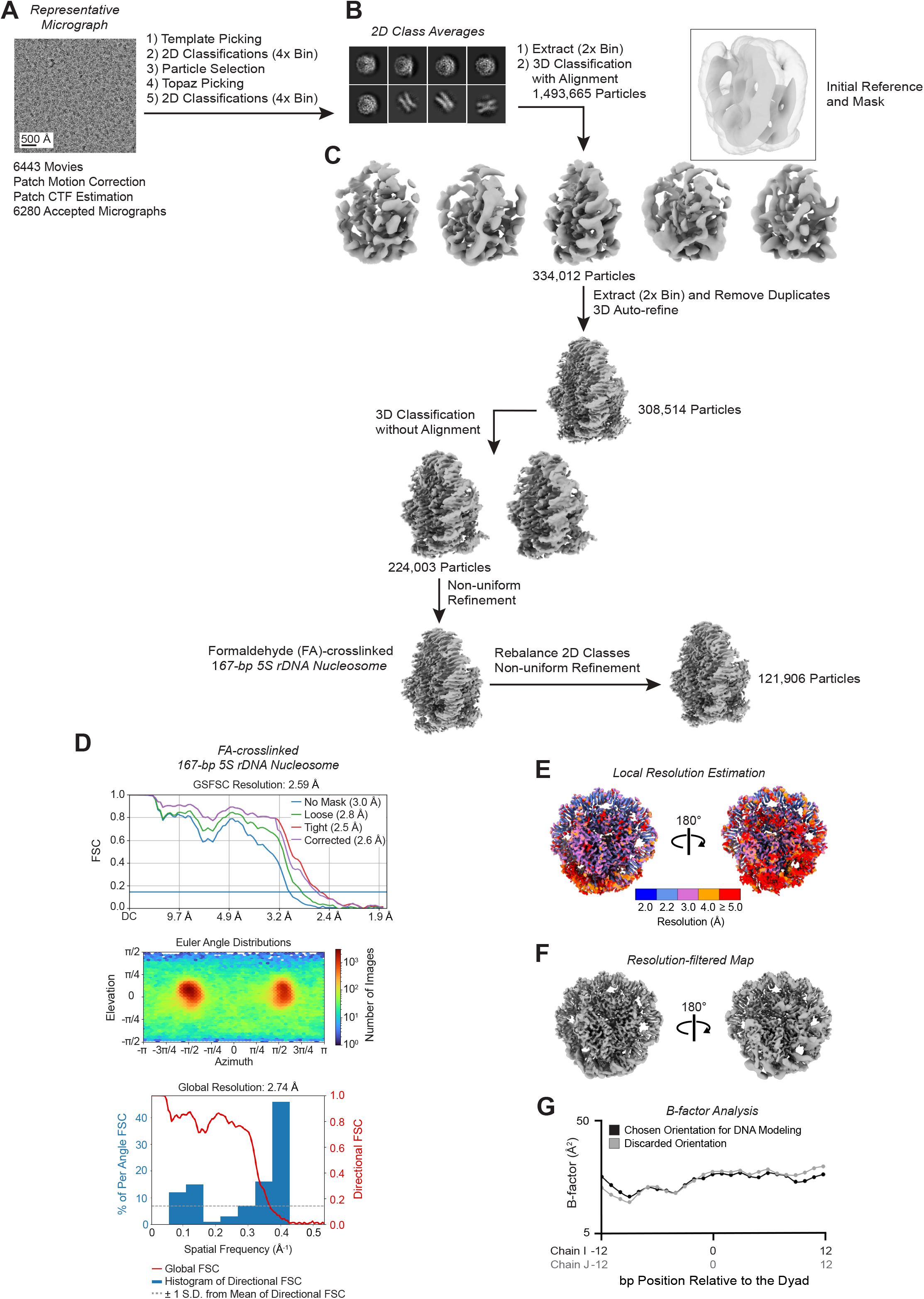
Image processing of the formaldehyde-crosslinked 167-bp 5S rDNA nucleosomes. (A) Representative micrograph, 10 Å low-pass filtered. (B) Selected 2D class averages. (C) 3D classifications and refinements. Note that the initial reference for 3D classifications with alignment presented one DNA end wrapped around the octamer but the 3D classification output volume displayed both ends flexible. (D) Fourier shell correlation (FSC), particle angle distribution, and histogram of directional FSC. (E) Local resolution estimation plotted onto the map. (F) Map filtered according to the local resolution estimation. (G) Average per-bp B factors plotted for the 25 bp refined in the two possible orientations.

**Supplementary Figure S19.**
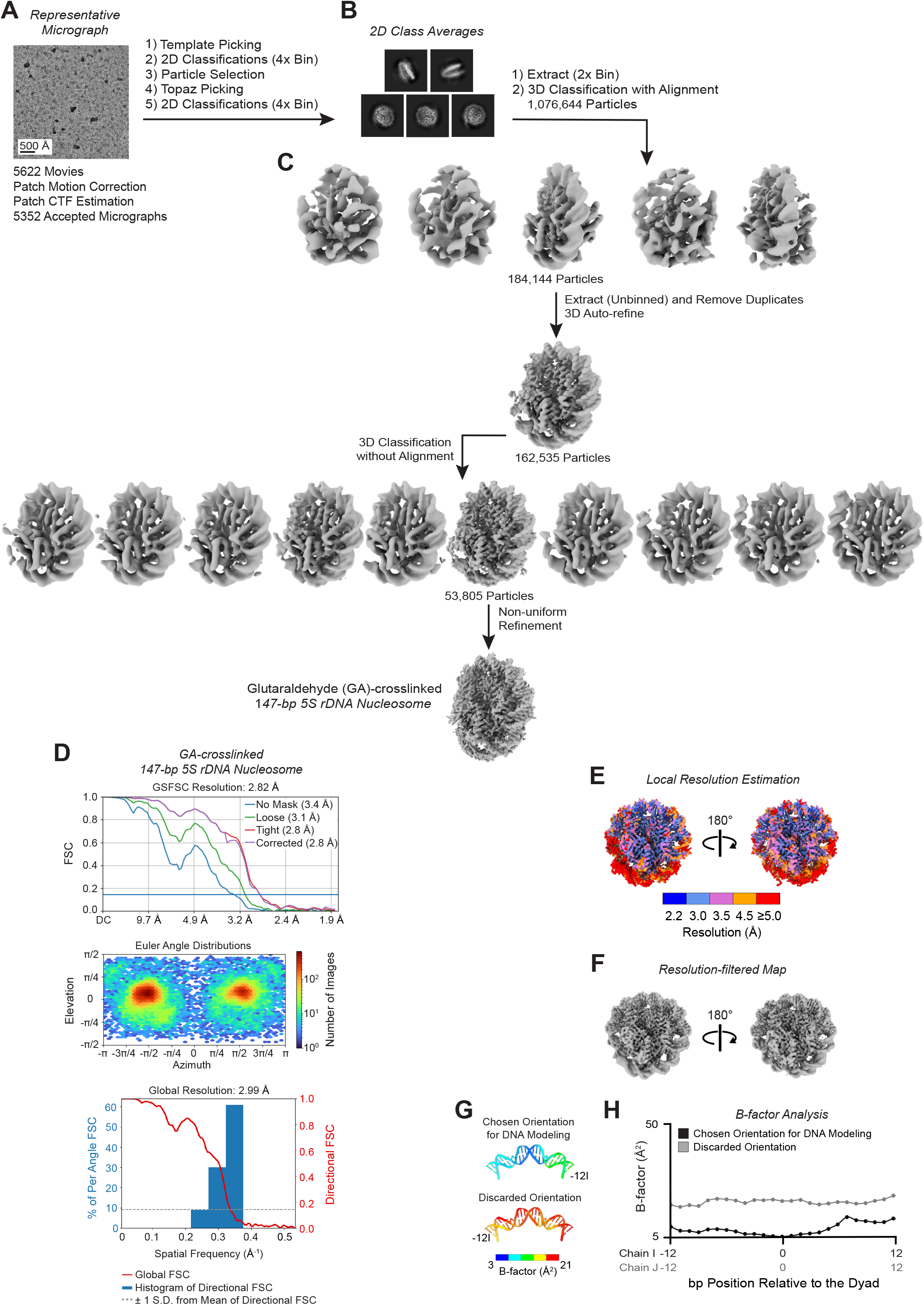
Image processing of the glutaraldehyde-crosslinked 147-bp 5S rDNA nucleosomes. (A) Representative micrograph, 10 Å low-pass filtered. (B) Selected 2D class averages. (C) 3D classifications and refinements. (D) Fourier shell correlation (FSC), particle angle distribution, and histogram of directional FSC. (E) Local resolution estimation plotted onto the map. (F) Map filtered according to the local resolution estimation. (G) DNA, real-space refined into the map, colored according to the per-bp B-factors (see Supplemental Materials and methods). Only the 25 bp of nucleosomal DNA centered on the dyad, which is the region of highest local resolution, are shown here. The DNA was refined in the two possible orientations (related through a 180° rotation along the dyad axis). We used the orientation with the lower B-factors to build the final atomic models. (H) Average per-bp B factors plotted for the 25 bp refined in the two possible orientations.

**Supplementary Figure S20.**
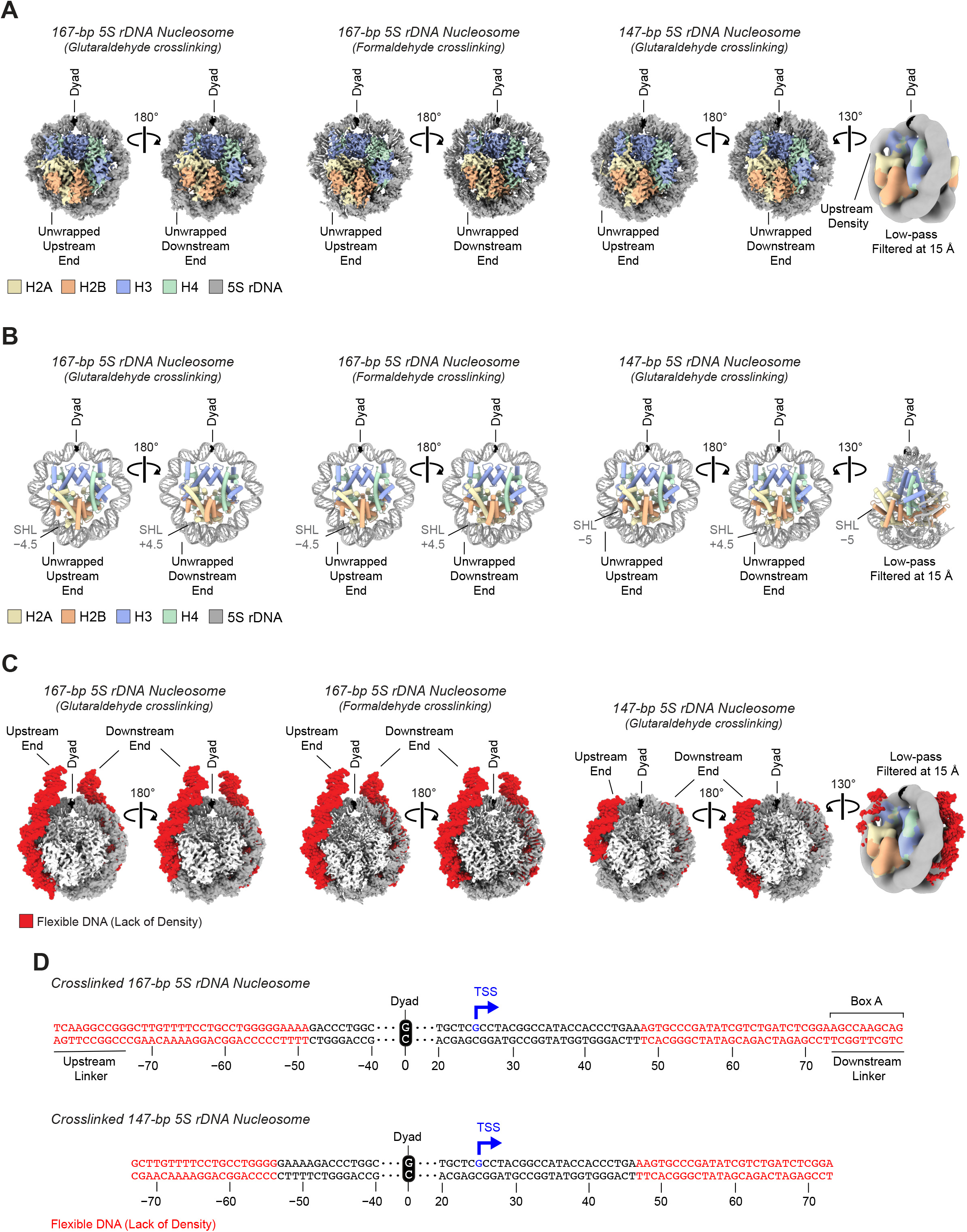
Crosslinking of the 167-bp and 147-bp 5S rDNA nucleosomes results in DNA unwrapping. (A) Cryo-EM density maps of the 5S rDNA nucleosomes crosslinked with either glutaraldehyde or formaldehyde. (B) Models of the crosslinked 5S rDNA nucleosomes. SHL: superhelical location. (C) Cryo-EM maps with the addition of red molecular models, which indicate the DNA segments that lack density (flexible DNA) in the crosslinked 5S rDNA nucleosomes. (D) Flexible DNA sequence in the crosslinked 5S rDNA nucleosomes. The DNA sequence in red type is not visible in the cryo-EM maps. The numbers indicate the bp position relative to the dyad. The 5S rDNA ends in panels A–C are labeled as upstream or downstream relative to the transcription start site (TSS). A low pass-filtered version of the maps (farthest right images in panels A–C) highlights density that is poorly resolved in the unfiltered map of the 147-bp 5S rDNA nucleosome. The full 167-bp and 147-bp 5S rDNA sequences are available in the Supplemental Materials and methods.

**Supplementary Figure S21.**
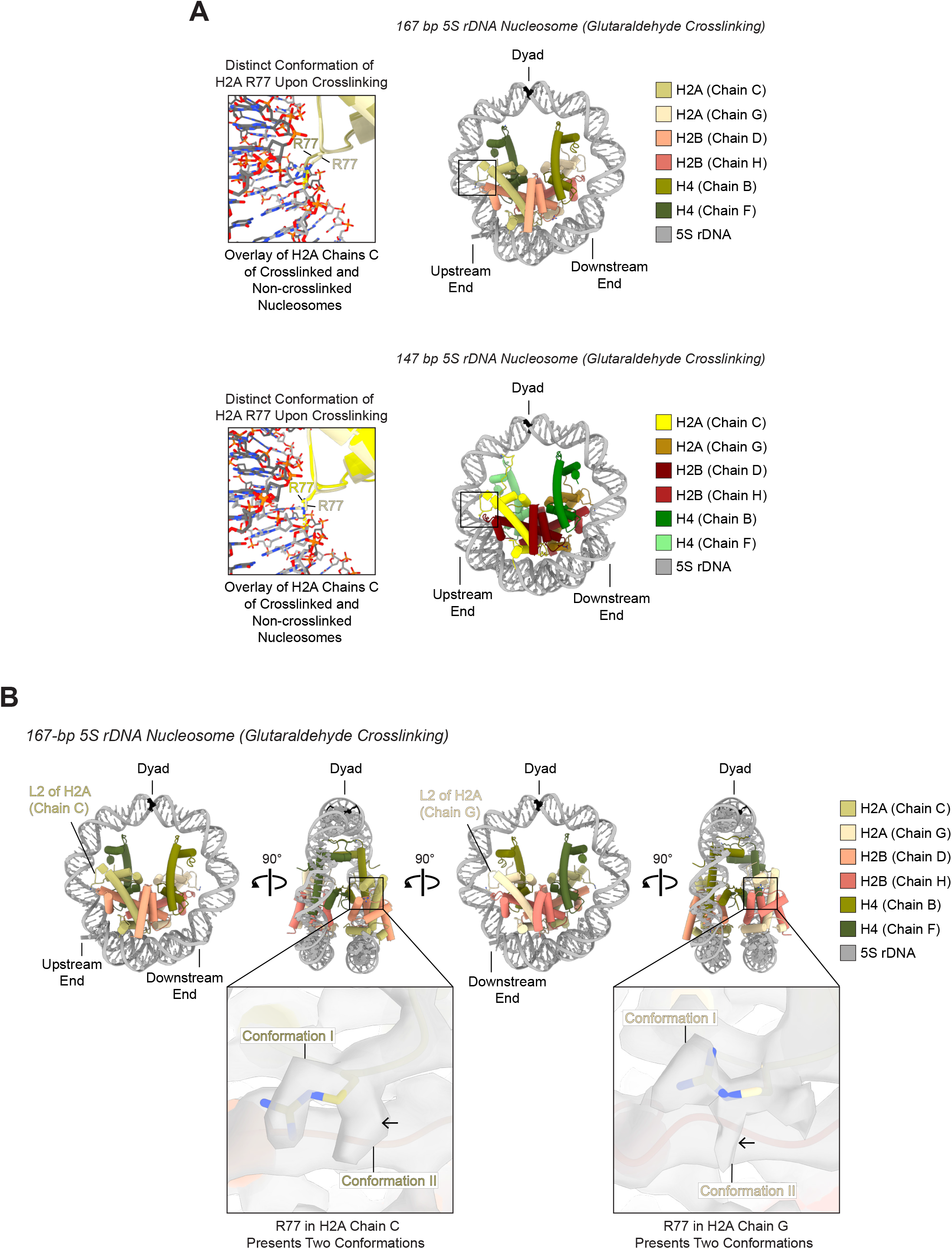
H2A R77 adopts two different conformations on each side of the glutaralde-hyde-crosslinked 167-bp 5S rDNA nucleosome. (A) Atomic models of the 167-bp and 147-bp 5S rDNA nucleosomes crosslinked with glutaraldehyde with close-up views of H2A R77. In the images on the left, the H2A chains C of crosslinked and non-crosslinked nucleosomes are overlaid to show that upon crosslinking H2A R77 adopts a conformation that would collide (depicted by yellow dashed lines) with a fully wrapped outer DNA turn. The H2A chains C of the non-crosslinked nucleosomes are the same as in Supplemental Fig. S15. Histone H3 is removed from the model for clarity. (B) The close-up views of the 167-bp 5S rDNA nucleosome crosslinked with glutaraldehyde show the cryo-EM density, in light gray, for the two conformations of H2A R77 on each of the two nucleosome sides, as well as the modeled H2A R77 (conformation I). The arrows indicate the unmodeled conformation (conformation II) of R77 in H2A chains C and G.

**Supplementary Figure S22.**
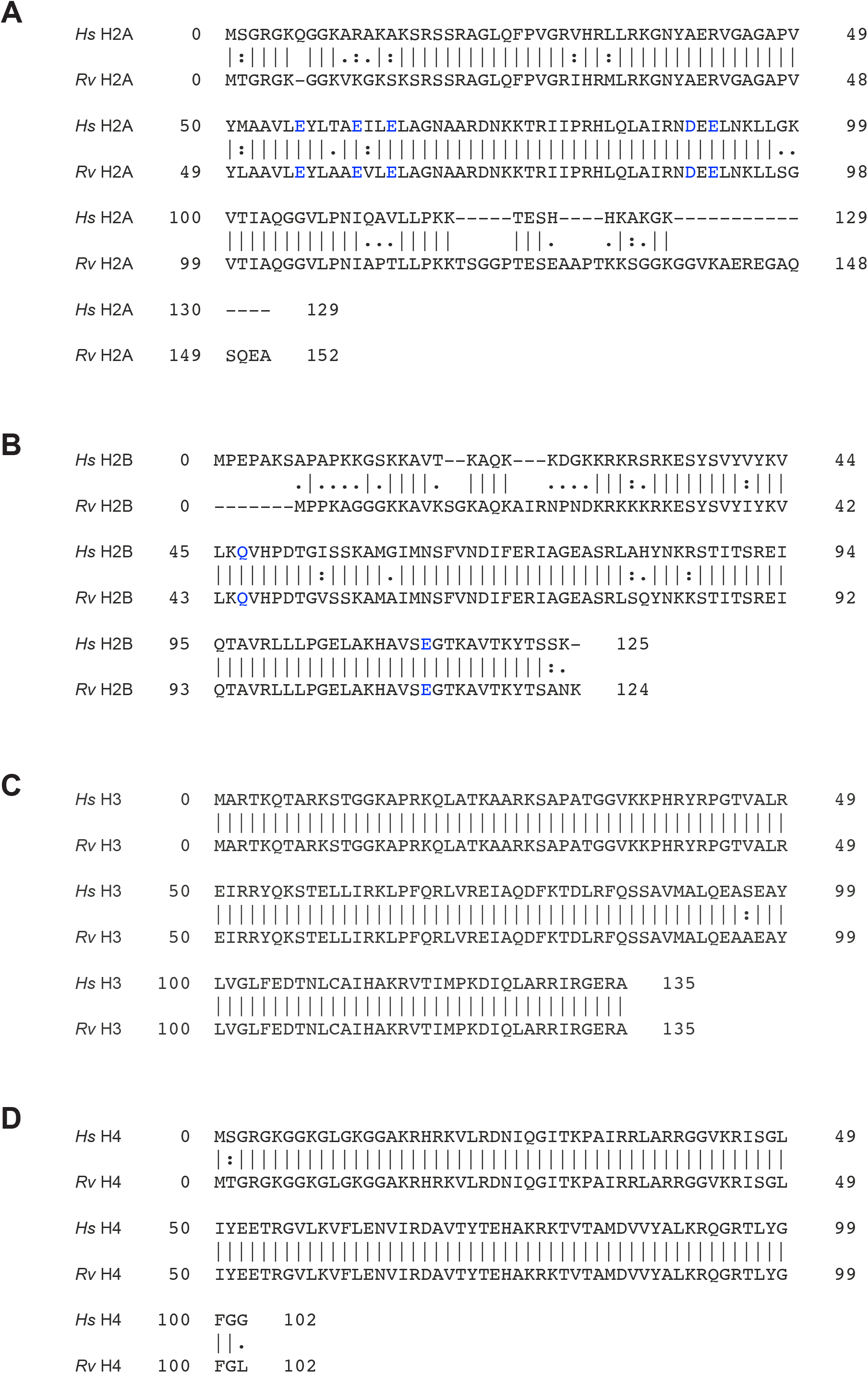
Sequence alignment of human and *R. varieornatus* core histone proteins. (A–D) The full-length sequences of human and *R. varieornatus* histones H2A, H2B, H3, and H4 were aligned using EMBOSS Needle (https://www.ebi.ac.uk/Tools/psa/emboss_needle/). The alignments reveal that the core histones, including the H2A and H2B amino acid residues involved in interactions with Dsup (indicated in blue type), are conserved between humans and *R. varieornatus*. The histone protein sequences from the *R. varieornatus* strain YOKOZUNA-1 (Hashimoto et al. 2016) were obtained using BLASTP (Altschul et al. 1997) against human histones, which were used to reconstitute nucleosomes in this study. To maintain consistency with the standard histone numbering nomenclature, the initiating methionine residues, which are typically removed from the histone proteins, are denoted as position zero (“0”) in each sequence.

**Supplementary Table S1.**
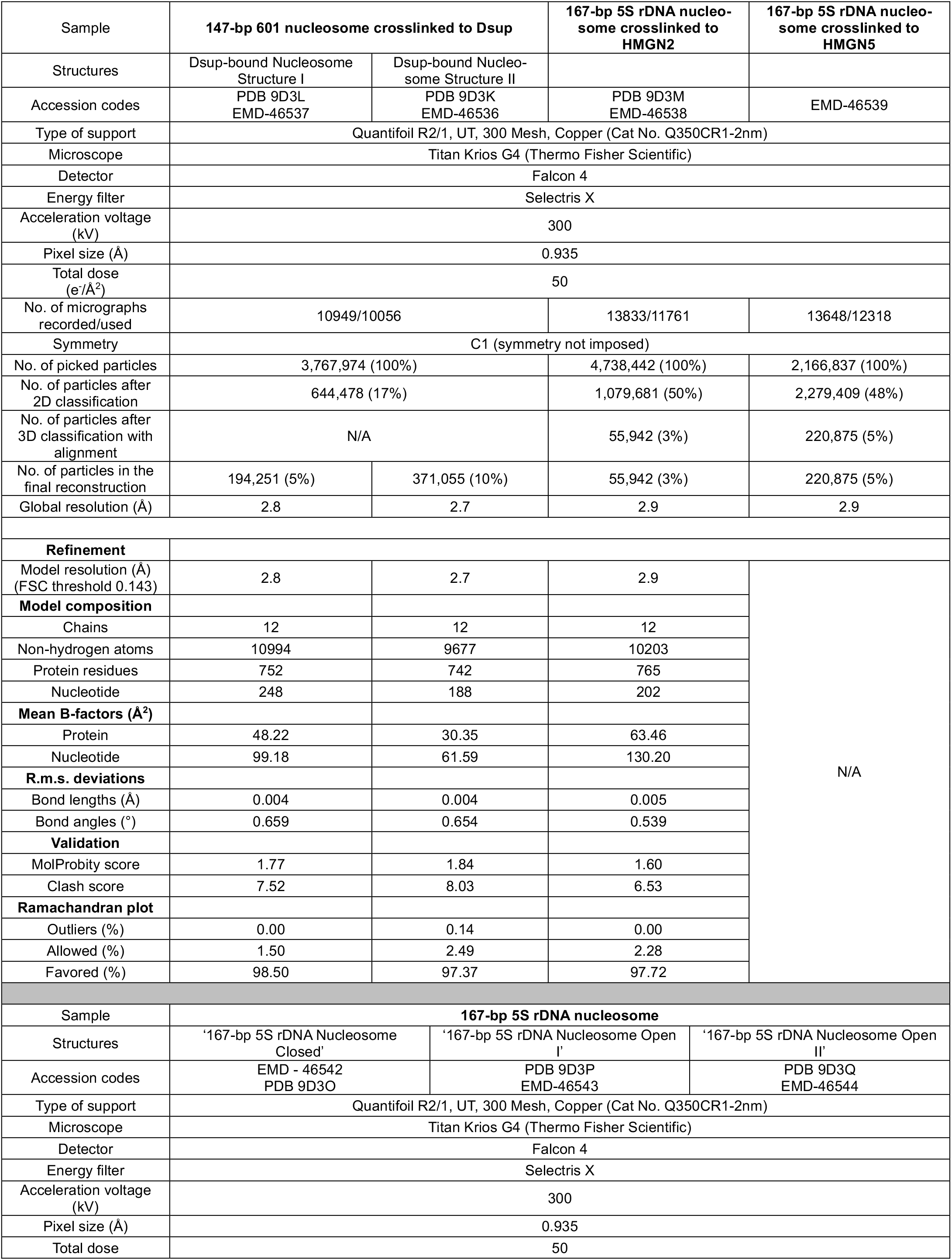
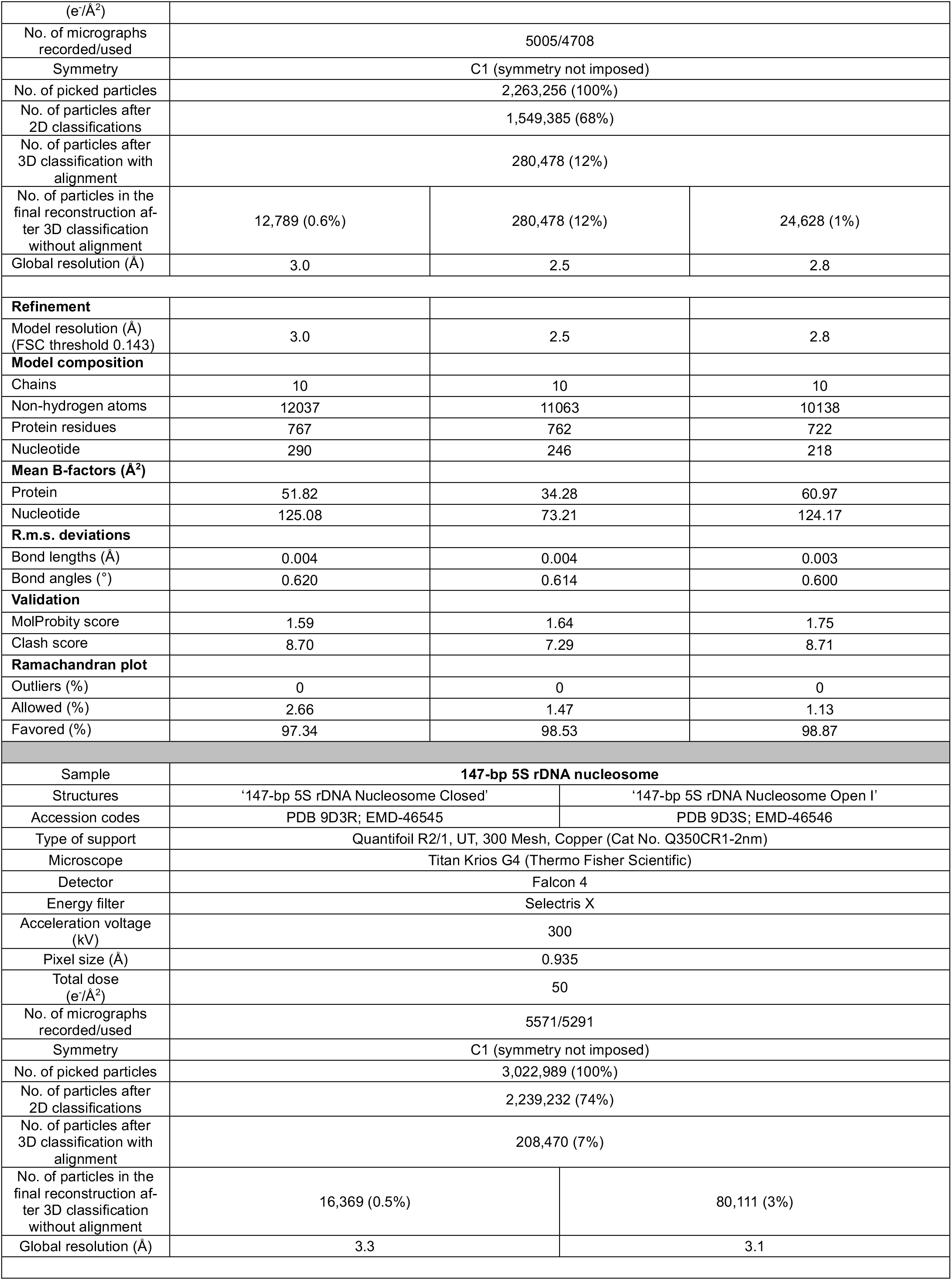
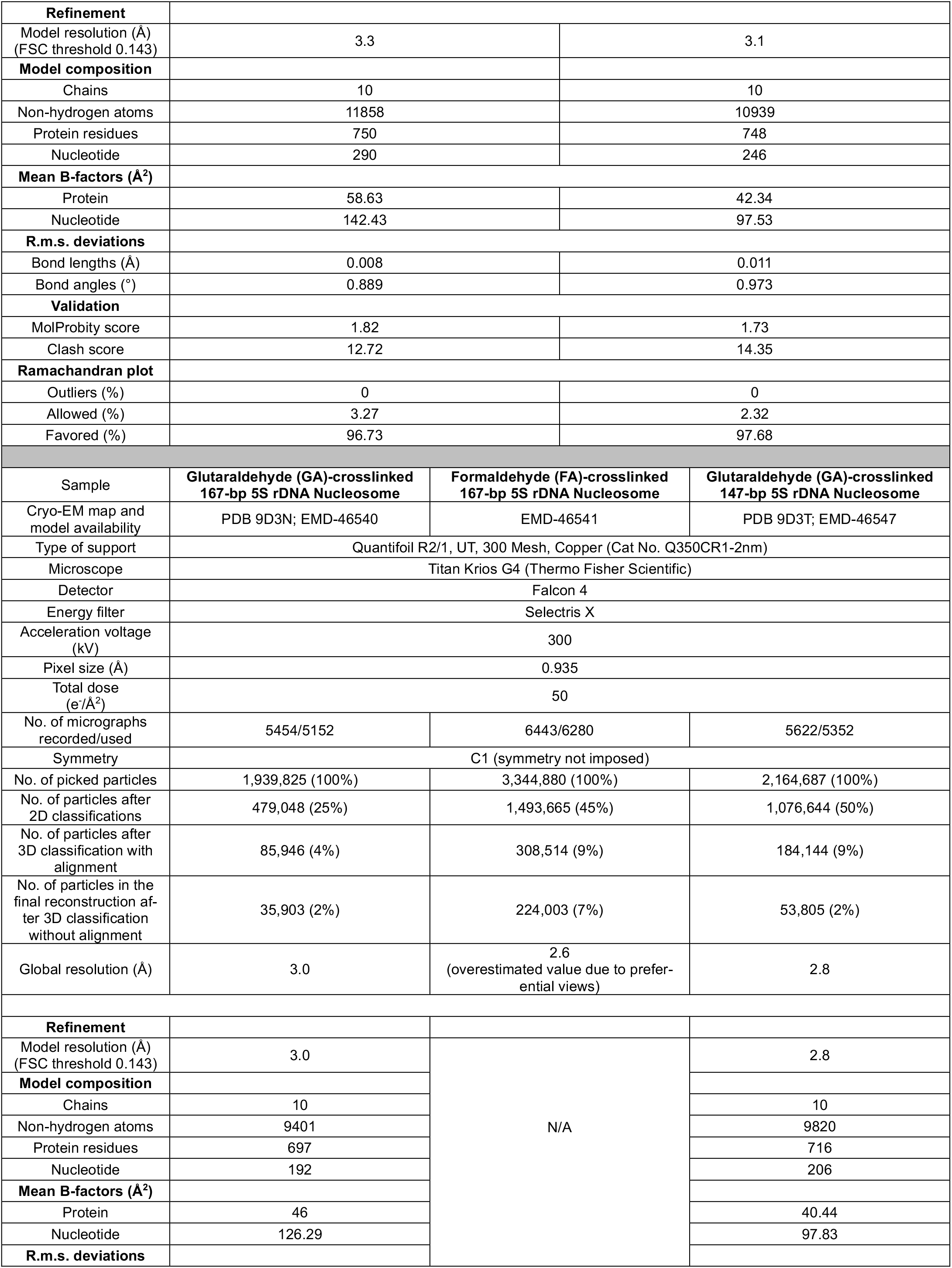
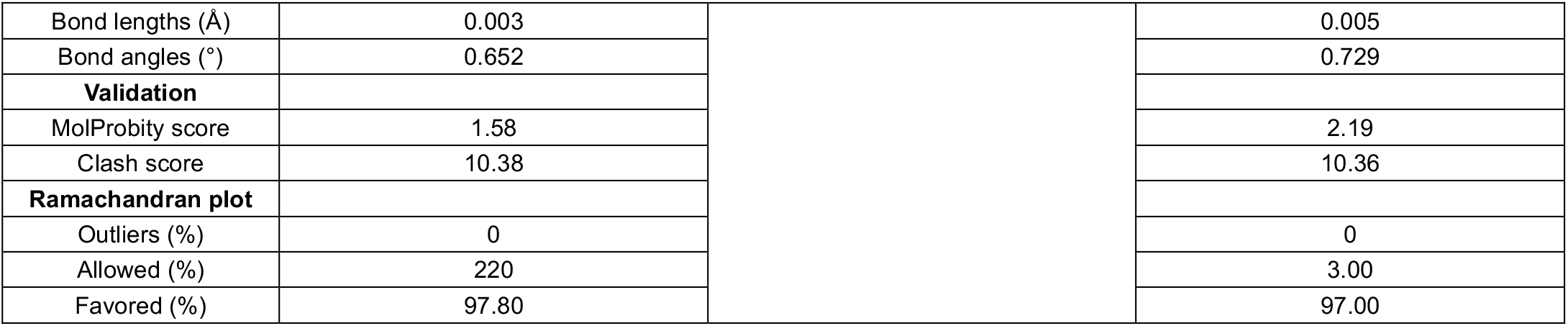
Cryo-EM data collection, refinement, and validation statistics.

| Sample | 147-bp 601 nucleosome crosslinked to Dsup |  | 167-bp 5S rDNA nucleosome crosslinked to HMGN2 | 167-bp 5S rDNA nucleosome crosslinked to HMGN5 |
| --- | --- | --- | --- | --- |
| Structures | Dsup-bound Nucleosome Structure I | Dsup-bound Nucleosome Structure II |  |  |
| Accession codes | PDB 9D3L<br>EMD-46537 | PDB 9D3K<br>EMD-46536 | PDB 9D3M<br>EMD-46538 | EMD-46539 |
| Type of support | Quantifoil R2/1, UT, 300 Mesh, Copper (Cat No. Q350CR1-2nm) |  |  |  |
| Microscope | Titan Krios G4 (Thermo Fisher Scientific) |  |  |  |
| Detector | Falcon 4 |  |  |  |
| Energy filter | Selectris X |  |  |  |
| Acceleration voltage (kV) | 300 |  |  |  |
| Pixel size (Å) | 0.935 |  |  |  |
| Total dose (e<sup>-</sup>/Å<sup>2</sup>) | 50 |  |  |  |
| No. of micrographs recorded/used | 10949/10056 |  | 13833/11761 | 13648/12318 |
| Symmetry | C1 (symmetry not imposed) |  |  |  |
| No. of picked particles | 3,767,974 (100%) |  | 4,738,442 (100%) | 2,166,837 (100%) |
| No. of particles after 2D classification | 644,478 (17%) |  | 1,079,681 (50%) | 2,279,409 (48%) |
| No. of particles after 3D classification with alignment | N/A |  | 55,942 (3%) | 220,875 (5%) |
| No. of particles in the final reconstruction | 194,251 (5%) | 371,055 (10%) | 55,942 (3%) | 220,875 (5%) |
| Global resolution (Å) | 2.8 | 2.7 | 2.9 | 2.9 |
| Refinement |  |  |  |  |
| Model resolution (Å) (FSC threshold 0.143) | 2.8 | 2.7 | 2.9 | N/A |
| Model composition |  |  |  |  |
| Chains | 12 | 12 | 12 |  |
| Non-hydrogen atoms | 10994 | 9677 | 10203 |  |
| Protein residues | 752 | 742 | 765 |  |
| Nucleotide | 248 | 188 | 202 |  |
| Mean B-factors (Å<sup>2</sup>) |  |  |  |  |
| Protein | 48.22 | 30.35 | 63.46 |  |
| Nucleotide | 99.18 | 61.59 | 130.20 |  |
| R.m.s. deviations |  |  |  |  |
| Bond lengths (Å) | 0.004 | 0.004 | 0.005 |  |
| Bond angles (°) | 0.659 | 0.654 | 0.539 |  |
| Validation |  |  |  |  |
| MolProbity score | 1.77 | 1.84 | 1.60 |  |
| Clash score | 7.52 | 8.03 | 6.53 |  |
| Ramachandran plot |  |  |  |  |
| Outliers (%) | 0.00 | 0.14 | 0.00 |  |
| Allowed (%) | 1.50 | 2.49 | 2.28 |  |
| Favored (%) | 98.50 | 97.37 | 97.72 |  |
| Sample | 167-bp 5S rDNA nucleosome |  |  |  |
| Structures | '167-bp 5S rDNA Nucleosome Closed' | '167-bp 5S rDNA Nucleosome Open I' | '167-bp 5S rDNA Nucleosome Open II' |  |
| Accession codes | EMD - 46542<br>PDB 9D3O | PDB 9D3P<br>EMD-46543 | PDB 9D3Q<br>EMD-46544 |  |
| Type of support | Quantifoil R2/1, UT, 300 Mesh, Copper (Cat No. Q350CR1-2nm) |  |  |  |
| Microscope | Titan Krios G4 (Thermo Fisher Scientific) |  |  |  |
| Detector | Falcon 4 |  |  |  |
| Energy filter | Selectris X |  |  |  |
| Acceleration voltage (kV) | 300 |  |  |  |
| Pixel size (Å) | 0.935 |  |  |  |
| Total dose | 50 |  |  |  |

|  |  |  |  |
| --- | --- | --- | --- |
| (e<sup>-</sup>/Å<sup>2</sup>) |  |  |  |
| No. of micrographs recorded/used | 5005/4708 |  |  |
| Symmetry | C1 (symmetry not imposed) |  |  |
| No. of picked particles | 2,263,256 (100%) |  |  |
| No. of particles after 2D classifications | 1,549,385 (68%) |  |  |
| No. of particles after 3D classification with alignment | 280,478 (12%) |  |  |
| No. of particles in the final reconstruction after 3D classification without alignment | 12,789 (0.6%) | 280,478 (12%) | 24,628 (1%) |
| Global resolution (Å) | 3.0 | 2.5 | 2.8 |
| Refinement |  |  |  |
| Model resolution (Å) (FSC threshold 0.143) | 3.0 | 2.5 | 2.8 |
| Model composition |  |  |  |
| Chains | 10 | 10 | 10 |
| Non-hydrogen atoms | 12037 | 11063 | 10138 |
| Protein residues | 767 | 762 | 722 |
| Nucleotide | 290 | 246 | 218 |
| Mean B-factors (Å<sup>2</sup>) |  |  |  |
| Protein | 51.82 | 34.28 | 60.97 |
| Nucleotide | 125.08 | 73.21 | 124.17 |
| R.m.s. deviations |  |  |  |
| Bond lengths (Å) | 0.004 | 0.004 | 0.003 |
| Bond angles (°) | 0.620 | 0.614 | 0.600 |
| Validation |  |  |  |
| MolProbity score | 1.59 | 1.64 | 1.75 |
| Clash score | 8.70 | 7.29 | 8.71 |
| Ramachandran plot |  |  |  |
| Outliers (%) | 0 | 0 | 0 |
| Allowed (%) | 2.66 | 1.47 | 1.13 |
| Favored (%) | 97.34 | 98.53 | 98.87 |
| Sample | 147-bp 5S rDNA nucleosome |  |  |
| Structures | ‘147-bp 5S rDNA Nucleosome Closed’ | ‘147-bp 5S rDNA Nucleosome Open I’ |  |
| Accession codes | PDB 9D3R; EMD-46545 | PDB 9D3S; EMD-46546 |  |
| Type of support | Quantifoil R2/1, UT, 300 Mesh, Copper (Cat No. Q350CR1-2nm) |  |  |
| Microscope | Titan Krios G4 (Thermo Fisher Scientific) |  |  |
| Detector | Falcon 4 |  |  |
| Energy filter | Selectris X |  |  |
| Acceleration voltage (kV) | 300 |  |  |
| Pixel size (Å) | 0.935 |  |  |
| Total dose (e<sup>-</sup>/Å<sup>2</sup>) | 50 |  |  |
| No. of micrographs recorded/used | 5571/5291 |  |  |
| Symmetry | C1 (symmetry not imposed) |  |  |
| No. of picked particles | 3,022,989 (100%) |  |  |
| No. of particles after 2D classifications | 2,239,232 (74%) |  |  |
| No. of particles after 3D classification with alignment | 208,470 (7%) |  |  |
| No. of particles in the final reconstruction after 3D classification without alignment | 16,369 (0.5%) | 80,111 (3%) |  |
| Global resolution (Å) | 3.3 | 3.1 |  |

|  |  |  |  |
| --- | --- | --- | --- |
| **Refinement** |  |  |  |
| Model resolution (Å)<br>(FSC threshold 0.143) | 3.3 |  | 3.1 |
| **Model composition** |  |  |  |
| Chains | 10 |  | 10 |
| Non-hydrogen atoms | 11858 |  | 10939 |
| Protein residues | 750 |  | 748 |
| Nucleotide | 290 |  | 246 |
| **Mean B-factors (Å<sup>2</sup>)** |  |  |  |
| Protein | 58.63 |  | 42.34 |
| Nucleotide | 142.43 |  | 97.53 |
| **R.m.s. deviations** |  |  |  |
| Bond lengths (Å) | 0.008 |  | 0.011 |
| Bond angles (°) | 0.889 |  | 0.973 |
| **Validation** |  |  |  |
| MolProbity score | 1.82 |  | 1.73 |
| Clash score | 12.72 |  | 14.35 |
| **Ramachandran plot** |  |  |  |
| Outliers (%) | 0 |  | 0 |
| Allowed (%) | 3.27 |  | 2.32 |
| Favored (%) | 96.73 |  | 97.68 |
| Sample | **Glutaraldehyde (GA)-crosslinked**<br>**167-bp 5S rDNA Nucleosome** | **Formaldehyde (FA)-crosslinked**<br>**167-bp 5S rDNA Nucleosome** | **Glutaraldehyde (GA)-crosslinked**<br>**147-bp 5S rDNA Nucleosome** |
| Cryo-EM map and<br>model availability | PDB 9D3N; EMD-46540 | EMD-46541 | PDB 9D3T; EMD-46547 |
| Type of support | Quantifoil R2/1, UT, 300 Mesh, Copper (Cat No. Q350CR1-2nm) |  |  |
| Microscope | Titan Krios G4 (Thermo Fisher Scientific) |  |  |
| Detector | Falcon 4 |  |  |
| Energy filter | Selectris X |  |  |
| Acceleration voltage<br>(kV) | 300 |  |  |
| Pixel size (Å) | 0.935 |  |  |
| Total dose<br>(e/Å<sup>2</sup>) | 50 |  |  |
| No. of micrographs<br>recorded/used | 5454/5152 | 6443/6280 | 5622/5352 |
| Symmetry | C1 (symmetry not imposed) |  |  |
| No. of picked particles | 1,939,825 (100%) | 3,344,880 (100%) | 2,164,687 (100%) |
| No. of particles after<br>2D classifications | 479,048 (25%) | 1,493,665 (45%) | 1,076,644 (50%) |
| No. of particles after<br>3D classification with<br>alignment | 85,946 (4%) | 308,514 (9%) | 184,144 (9%) |
| No. of particles in the<br>final reconstruction af-<br>ter 3D classification<br>without alignment | 35,903 (2%) | 224,003 (7%) | 53,805 (2%) |
| Global resolution (Å) | 3.0 | 2.6<br>(overestimated value due to prefer-<br>ential views) | 2.8 |
| **Refinement** |  | N/A |  |
| Model resolution (Å)<br>(FSC threshold 0.143) | 3.0 |  | 2.8 |
| **Model composition** |  |  |  |
| Chains | 10 |  | 10 |
| Non-hydrogen atoms | 9401 |  | 9820 |
| Protein residues | 697 |  | 716 |
| Nucleotide | 192 |  | 206 |
| **Mean B-factors (Å<sup>2</sup>)** |  |  |  |
| Protein | 46 |  | 40.44 |
| Nucleotide | 126.29 |  | 97.83 |
| **R.m.s. deviations** |  |  |  |

Supplemental Table S1. Cryo-EM data collection, refinement, and validation statistics.
|  |  |  |  |
| --- | --- | --- | --- |
| Bond lengths (Å) | 0.003 |  | 0.005 |
| Bond angles (°) | 0.652 |  | 0.729 |
| Validation |  |  |  |
| MolProbity score | 1.58 |  | 2.19 |
| Clash score | 10.38 |  | 10.36 |
| Ramachandran plot |  |  |  |
| Outliers (%) | 0 |  | 0 |
| Allowed (%) | 220 |  | 3.00 |
| Favored (%) | 97.80 |  | 97.00 |

**Supplemental Movie S1. 167-bp 5s rDNA nucleosome dynamics: downstream DNA opening**.

**Supplemental Movie S2. 167-bp 5s rDNA nucleosome dynamics: upstream DNA opening**.

**Supplemental Movie S3. 3D variability analysis of the crosslinked Dsup-bound nucleosome after map refinement using an initial reference without one DNA end**.

**Supplemental Movie S4. 3D variability analysis of the crosslinked Dsup-bound nucleosome after map refinement using an initial reference without both DNA ends**.

## Supplemental Materials and methods

### Purification of recombinant human histones and reconstitution of histone octamers

The coding sequences of the four human core histones were each individually cloned into the pET11a vector (Novagen). Each histone was synthesized in *Escherichia coli* BL21(DE3) cells (Novagen) and purified from inclusion bodies by the method of Luger et al. (1999) with some modifications as follows. After induction with IPTG at a final concentration of 0.4 mM at 37 °C for 2 h, cells (1 g) were resuspended in 25 mL of cold buffer W (50 mM Tris-HCl, pH 7.5, 0.1 M NaCl, 1 mM EDTA, 100 µg/mL lysozyme, 1 mM benzamidine, 5 mM 2-mercaptoethanol, and 0.25 mM PMSF) and subjected to two freeze-thaw cycles. Next, the cell suspension was centrifuged (14,000 rpm for 15 min at 4 °C; Fiberlite F21S-8×50y rotor), and the resulting pellet was resuspended in 25 mL of cold TW buffer [50 mM Tris-HCl, pH 7.5, 0.1 M NaCl, 1 mM EDTA, 1% (v/v) Triton X-100, 1 mM benzamidine, 5 mM 2-mercaptoethanol, and 0.25 mM PMSF] and dispersed by using a glass Wheaton Dounce homogenizer with a loose B pestle. Then, the cells were lysed on ice by 10 sonication cycles (each for 15 s ON and 15 s OFF, 20% output; Branson Sonifier 450). The pellet containing the inclusion bodies was recovered by centrifugation (14,000 rpm for 15 min at 4 °C; Fiberlite F21S-8×50y rotor) and washed an additional two times with cold TW buffer followed by three washes with cold buffer W. The pellet was minced in 0.3 mL of DMSO, incubated for 30 min at 22 °C, suspended in 5 mL of buffer G (20 mM Tris-HCl, pH 7.5, 7 M guanidine-HCl, and 10 mM DTT), and incubated at 22 °C for 1 h on a nutator. The soluble fraction containing the histone was recovered by centrifugation (14,000 rpm for 15 min at 20 °C; Fiberlite F21S-8×50y rotor) and dialyzed (molecular weight cutoff: 3.5 kDa) against three changes (each at 4 °C for 2 h) of buffer U (10 mM Tris-HCl, pH 7.5, 1 mM EDTA, 7 M urea, 5 mM 2-mercaptoethanol, and 0.25 mM PMSF) containing 0.2 M NaCl. After dialysis, the insoluble material was removed by centrifugation (18,000 rpm for 10 min at 10 °C; Fiberlite F21S-8×50y rotor), and the histones were purified from the supernatant by tandem Q Sepharose High Performance followed by SP Sepharose High Performance ion-exchange chromatography. Briefly, after sample loading, the Q Sepharose High Performance column (5 mL packed resin) was removed, and the SP Sepharose High Performance column (5 mL packed resin) was washed with 15 mL of buffer U containing 0.2 M NaCl followed by 5 mL of buffer U containing 0.25 M NaCl. Then, the histones were eluted with a linear gradient of 0.25 M to 0.5 M NaCl in buffer U over 50 mL total volume. The peak fractions containing the most highly purified histones were pooled and dialyzed (molecular weight cutoff: 3.5 kDa) against three changes of 2 L of cold water containing 5 mM 2-mercaptoethanol and 0.2 mM PMSF (the first and the last dialysis steps for 2 h, and the second step overnight at 4 °C). After dialysis, the insoluble materials were removed by centrifugation, and the histones were lyophilized to dryness (medium drying rate; Savant Speed Vac Plus) and stored at −80 °C. The molecular mass of each purified histone protein was determined by mass spectrometry (Molecular Mass Spectrometry Facility, UCSD). This analysis confirmed the integrity of the histones and showed that all four histones lacked the N-terminal initiating methionine residue. The numbering of the histone residues throughout the figures and manuscript begins at the first residue following the initiating methionine.

### The sequences of the human core histone proteins used in this study

are as follows. Histone H2A2A (NCBI Reference Sequence NP_003500.1; UniProt Q6FI13): MSGRGKQGGKARAKAKSRSSRAGLQFPVGRVHR-LLRKGNYAERVGA-GAPVYMAAVLEYLTAEILELAGNAARDNKKTRIIPRHLQLAIRNDEELNKLLGKVTIAQGGVLPNIQAVLLPKKTESHHKAKGK; histone H2B1C (NCBI Reference Sequence NP_001368918.1; UniProt P62807): MPEPAKSAPAPKKGSKKAVTKAQKKDGKKRKRS-RKESYSVYVYKVLKQVHPDT-GISSKAMGIMNSFVNDIFERIAGEASRLAHYNKRSTITSREIQTAVRLLLPGELAKHAVSEGTKAVTKYTSSK; histone H3.2 (NCBI Reference Sequence NP_001005464.1; UniProt Q71DI3): MARTKQTARKSTGGKAPRKQLATKAARKSAPATGGVKKPHRYRPGT-VALREIRRYQK-STELLIRKLPFQRLVREIAQDFKTDLRFQSSAVMALQEASEAYLVGLFEDTNLCAIHAKRVTIMPKDIQLARRIRGERA; histone H4 (NCBI Reference Sequence NP_001029249.1; UniProt P62805): MSGRGKGGKGLGKGGAKRHRKVLRDNIQGITKPAIRR-LARRGGVKRISGLIYEET-RGVLKVFLENVIRDAVTYTEHAKRKTVTAMDVVYALKRQGRTLYGFG G.

To reconstitute histone octamers, each lyophilized histone was dissolved in buffer G at 22 °C for 1 h. The absorbance at 280 nm of each histone sample was measured with a spectrophotometer (Nanodrop One; Thermo Scientific), and the histone concentration was determined by using the following molar extinction coefficients at 280 nm and molecular masses: H2A2A, 4470 M^-1^ cm^-1^, 14,095 g/mol; H2B1C, 7450 M^-1^ cm^-1^, 13,906 g/mol; H3.2, 4470 M^-1^ cm^-1^, 15,388 g/mol; H4, 5960 M^-1^ cm^-1^, 11,367 g/mol. Then, a histone mix was prepared by combining an equimolar ratio of H3 and H4 with a 25% molar excess of both H2A and H2B (*i*.*e*., molar ratio 1.0:1.0:1.25:1.25 of H3:H4:H2A:H2B). The final protein concentration was adjusted to 1 mg/mL with buffer G, and the mixture (3 mL maximum volume) was dialyzed (molecular weight cutoff: 3.5 kDa) against three changes of 1 L of buffer R (10 mM Tris-HCl, pH 7.5, 2 M NaCl, 1 mM EDTA, and 5 mM 2-mercaptoethanol) at 4 °C (the first and the last dialysis steps for 2 h each, and the second step overnight). The insoluble material was removed by centrifugation at 13,200 rpm for 10 min at 4 °C (Eppendorf 5415R), and the supernatant was concentrated by using a protein concentrator (molecular weight cutoff: 50 kDa). The concentrated sample was centrifuged at 13,200 rpm for 10 min at 4 °C (Eppendorf 5415R), and the supernatant (about 0.24 mL) was loaded onto a Superose 12 HR 10/30 size exclusion column (24 mL packed resin; GE Healthcare). The histone octamers were eluted with 24 mL of buffer R (fraction size: 0.25 mL). The peak fractions containing equimolar amounts of each histone were pooled and then dialyzed against three changes of storage buffer [10 mM Hepes-K^+^, pH 7.6, 1 mM EDTA, 10 mM KCl, 10% (v/v) glycerol, and 1 mM DTT] at 4 °C (the first and the last dialysis steps for 2 h each, and the second step overnight). The histones were frozen in liquid nitrogen and stored at −80 °C.

### Purification of recombinant Dsup and recombinant HMGN proteins

A recombinant His6- and FLAG-tagged version of *R. varieornatus* Dsup was purified by the nondenaturing method described in Chavez et al. (2019).

The coding sequence of human HMGN2 protein was codon optimized for expression in *E. coli* and cloned into vector pET11a (Novagen). BL21(DE3) cells were transformed with the recombinant DNA construct, and protein synthesis was induced by the addition of IPTG to a final concentration of 0.4 mM at 30 °C for 1.5 h. Then, HMGN2 was purified by cation-exchange chromatography as described in Paranjape et al. (1995) with some modifications. Briefly, 1 g of bacterial cells was resuspended in 6 mL of cold buffer L (50 mM Hepes-K^+^, pH 7.6, 0.1 M NaCl, 1 mM benzamidine, 1 mM DTT, 4 µg/mL leupeptin, 4 µg/mL aprotinin, 1 µg/mL pepstatin, and 0.2 mM PMSF) and then lysed by sonication on ice (10 cycles 15 s ON/ 15 s OFF, 20% output; Branson Sonifier 450). The insoluble fraction was recovered by centrifugation (13,000 rpm for 10 min at 4 °C; Fiberlite F21S-8×50y rotor). The pellet containing HMGN2 was resuspended in 4 mL of cold buffer L, and the protein was extracted by the addition of 12 M HCl to a final concentration of 0.77 M. After incubation at 4 °C for 1 h on a rotating wheel, the insoluble material was removed by centrifugation (13,000 rpm for 10 min at 4 °C; Fiberlite F21S-8×50y rotor), and the supernatant containing HMGN2 was dialyzed (molecular weight cutoff: 3.5 kDa) against two changes of 2 L of buffer H (50 mM Hepes-K^+^, pH 7.6, 0.1 M NaCl, 1 mM benzamidine, 1 mM DTT, and 0.2 mM PMSF), with each dialysis step at 4 °C for 2 h. After dialysis, the insoluble material was removed by centrifugation (18,000 rpm for 10 min at 4 °C; Fiberlite F21S-8×50y rotor), and the supernatant containing HMGN2 was loaded onto a CM Sepharose Fast Flow column (20 mL packed resin; column: XK16/20 GE Healthcare). The column was washed with 80 mL of buffer H, and then the protein was eluted with a 0.15 M to 0.35 M NaCl gradient in buffer H over 60 mL followed by a 0.35 M to 0.55 M NaCl gradient in buffer H over 100 mL. The peak fractions containing highly pure HMGN2 were pooled and dialyzed against two changes of buffer S [10 mM Hepes-K^+^, pH 7.6, 10 mM KCl, 0.5 mM EGTA, 1.5 mM MgCl_2_, 10% (v/v) glycerol, 1 mM DTT, 10 mM glycerol 2-phosphate, 1 mM benzamidine, and 0.2 mM PMSF] at 4 °C (the first dialysis step for 2 h and the second dialysis step overnight). The protein was frozen in liquid nitrogen and stored at −80 °C. The molecular mass of the human HMGN2 protein was determined by mass spectrometry (Molecular Mass Spectrometry Facility, UCSD). This analysis confirmed the integrity of the protein and showed that HMGN2 lacks the N-terminal initiating methionine residue. The numbering of the HMGN2 residues throughout the figures and manuscript begins at the first residue following the initiating methionine. The human HMGN2 protein sequence is as follows (NCBI Reference Sequence NP_005508.1; UniProt P05204): MPKRKAEGDAKGDKAKVK-DEPQRRSARLSAKPAPPKPEPKPKKA-PAKKGEKVPKGKKGKADAGKEGNNPAENGDAKTDQAQKAEGAGDA K.

The coding sequence of the human HMGN5 protein containing His6 (N-terminal) and FLAG (C-terminal) tags was codon optimized for expression in *E. coli* and cloned into the pET21b vector (Novagen). The His6-FLAG tagged HMGN5 protein was synthesized in Rosetta(DE3)pLysS cells (Novagen) and purified by Ni-NTA affinity chromatography as described for Dsup in Chavez et al. (2019) with the following modifications. First, the lysis buffer was substituted with buffer HL [50 mM Hepes-Na^+^, pH 7.8, 0.1 mM EDTA, 0.1 M NaCl, 0.01% (v/v) NP-40, 5% (v/v) glycerol, 10 mM 2-mercaptoethanol, 1 µg/mL pepstatin, 1 µg/mL leupeptin, 1 mM benzamidine, 0.5 mM PMSF, 300 µg/mL lysozyme, and 100 µg/mL Pefabloc (4-(2-aminoethyl)-benzene-1-sulfonyl fluoride)]. Second, after sonication in buffer HL, the protein was extracted from the soluble material by the addition of 12 M HCl to a final concentration of 0.75 M for 30 min on ice. Then, the supernatant containing the protein was recovered by centrifugation (18,000 rpm for 15 min at 4 °C; Fiberlite F21S-8×50y rotor). The solution was neutralized by the addition of 5 M NaOH, and 1 M Tris-HCl, pH 8.0, and 1 M imidazole were added to 50 mM and 20 mM final concentrations, respectively, before loading onto a Ni-NTA column. The human HMGN5 protein sequence is as follows (NCBI Reference Sequence NP_110390.1; UniProt P82970): MPKRKAAGQGDMRQEPKRRSARLSAM-LVPVTPEVKPKRTSSSRKMKTKSDMMEENIDTSAQA-VAETKQEAVVEEDYNENAKNGEAKITEAPASEKEIVEVKEENIEDATEKGGEKKEAVAAEVKNEEEDQKEDEEDQNEEKGEAGKEDK-DEKGEEDGKEDKNGNEKGEDAKEKEDGKKGEDGKGNGEDGKEKGE DEKEEEDRKETGDGKENEDGKEKGDKKEGKDVKVKEDEKEREDGK-EDEGGNEEEAGKEKEDLKEEEEGKEEDEIKEDDGKKEEPQSIV.

### Reconstitution of nucleosomes as well as Dsup-nucleosome and HMGN-nucleosome complexes

Nucleosomes (mononucleosomes) containing recombinant human histones octamers and 147 bp or 167 bp of *Xenopus borealis* 5S rDNA, or 147 bp of the 601 DNA sequence were reconstituted by step-wise salt dialysis as described in Chavez et al. (2019) with the following modifications. Briefly, the DNA was combined with the histone octamers in buffer M [25 mM Hepes-K^+^, pH 7.6, 0.1 mM EDTA, 0.01% (v/v) NP-40, and 1 M NaCl]. The histone-DNA mixture was incubated for 30 min on ice and then dialyzed (molecular weight cutoff: 3.5 kDa) against three changes (each at 22 °C for 2.5 h) of buffer HE (25 mM Hepes-K^+^, pH 7.6, and 0.1 mM EDTA) containing decreasing concentrations of NaCl (0.8 M NaCl, 0.6 M NaCl, and 50 mM NaCl). Then, the samples were heated at 58 °C for 10 min to facilitate the most stable nucleosome position, and the quality of the nucleosomes was assessed by native (nondenaturing) 4.5% polyacrylamide gel electrophoresis. The reconstituted nucleosomes were stored at 4 °C prior to sample preparation for cryo-EM studies. The 147-bp 601-DNA sequence is as follows: CTGGAGAATCCCGGTGCCGAGGCCGCTCAATTGGTCGTAGACAGCTCTAGCACCGCTTAAAC-GCACGTACGCGCTGTCCCCCGCGTTTTAACCGCCAAGGGGATTACTCCCTAGTCTCCAGGCACGTGTCAGATATATACATCCTGT. The 147-bp *Xenopus borealis* somatic 5S rDNA gene fragment (Peterson et al. 1980; Rhodes et al. 1985) is the central 147-bp nucleosome positioning sequence that was precisely mapped in Fei et al. (2018). The sequence of the 167-bp *Xenopus borealis* 5S rDNA fragment (with the central 147-bp positioning sequence underlined) is as follows: TCAAGGCCGG<u>GCTT-</u> <u>GTTTTCCTGCCTGGGGGAAAAGACCCTGGCATGGGGAG-</u> <u>GAGCTGGGCCCCCCCCAGAAGGCAGCACAAGGGGAGGAAAAGTCA</u> <u>GCCTTGTGCTCGCCTACGGCCATACCACCCTGAAAGTGCCCGA-</u> <u>TATCGTCTGATCTCGGA</u>AGCCAAGCAG.

To reconstitute the protein-nucleosome complexes (2.1 µM final concentration), the purified recombinant Dsup, HMGN2, or HMGN5 proteins were combined with nucleosomes in binding buffer [25 mM Hepes-K^+^, pH 7.6, 0.1 mM EDTA, 90 mM KCl, 1.8% (v/v) glycerol, and 0.01% (v/v) NP-40] and incubated at 27 °C for 1 h. The molar ratios of nucleosome:Dsup, nucleosome:HMGN2, and nucleosome:HMGN5 in the binding reactions were 1:3, 1:3, and 1:2, respectively. In addition, control samples of nucleosomes (2.1 µM final concentration) that did not contain Dsup or the HMGN proteins were processed in parallel. The formation of the protein-nucleosome complexes was assessed by analyzing a small portion (1 pmol) of the samples by 4.5% polyacrylamide native gel electrophoresis.

### Grid preparation for cryo-EM analysis

Holey grids with continuous carbon on top (Quantifoil® R 2/1, UT, 300 Mesh, Cu or Quantifoil® R 2/1, UT, 200 Mesh, Cu (Electron Microscopy Sciences) were glow-discharged for 30 s at 20-30 mA. 2.6 µL to 3 µL of sample was applied to the grids and flash-frozen using a Vitrobot Mark IV (FEI; 95% humidity, 4 °C, blot force 4, blot time 4). Non-crosslinked samples were used at concentrations of 0.5-0.6 µM, whereas HMGN2 or Dsup crosslinked to nucleosomes were at 1.8 µM or 0.9 µM, respectively. All samples were kept on ice prior to vitrification.

### Cryo-EM data collection

Data were collected on the in-house Titan Krios G4 (Thermo Fisher Scientific) operated at 300 kV and equipped with a Falcon 4 direct electron detector (Thermo Fisher Scientific) and a Selectris X energy filter (Thermo Fisher Scientific). A nominal magnification of 130,000× (object pixel size of 0.935 Å), a total dose of 50 e^-^/Å^2^ and an applied defocuses range of −0.6 to −2.2 µm were used.

### Cryo-EM image processing

In CryoSPARC Live, on-the-fly Patch Motion Correction and Patch CTF Estimation jobs were used to apply the gain reference, align and estimate the CTF of the collected movie frames (Punjani et al. 2017). Micrographs with a CTF fit worse than 3 Å were always discarded. Depending on the dataset, the ‘Relative ice thickness’ (greater than 1.07) and ‘Total motion’ (greater than 20 pixels) parameters were also used to remove micrographs, as well as manual inspection. In CryoSPARC (Punjani et al. 2017), template picking followed of rounds of 2D classifications with 4x binned particles allowed the selection of ‘good’ particles, which were used for training Topaz (Bepler et al. 2019). Several rounds of 2D classification and particle curation culminated in the selection of a particle stack. For free and HMGN2-bound 5S nucleosomes, this particle stack was re-extracted at 1.87 Å/pix (2x binned). pyem/csparc2star.py (https://doi.org/10.5281/zenodo.3576630) was used to convert the CryoSPARC readable particle file to RELION’s readable format (.star) and 3D classifications with alignment were performed in RELION-3.1 or RELION-4.0 (Scheres 2012; Scheres 2016). A soft masked, 60 Å lowpass filtered initial reference was used, as well as a 113 to 120 Å circular mask around each particle. Initially, 25 iterations with exhaustive angular searches (angular sampling interval 7.5°, offset search range 5 pix, offset search step 1 pix) and regularization parameter T = 4 were run; then the 3D classification was continued 10 to 25 extra iterations with narrower angular searches (angular sampling interval 3.7°, offset search range 3 to 4 pix, offset search step 0.8 to 1 pix). The output class with best nucleosome features was 3D refined in RELION (Scheres, 2012) or CryoSPARC (Punjani et al. 2020), and aligned particles were 3D classified without alignment in RELION-3.1 or RELION-4.0. The initial reference was low-pass filtered at 10 to 15 Å, and it was softly masked; 2 to 10 classes were used. In the case of the HMGN2-bound 167-bp 5S rDNA nucleosome, 3D classification without alignment was not performed. Instead, a second round of 3D classification with alignment was run without masking the initial reference. After 3D classifications, particles from selected classes were imported to CryoSPARC and subjected to Non-uniform refinement in CryoSPARC (Punjani et al. 2020) (per-micrograph CTF refinement was always used; sometimes, enabling ‘Enforce non-negativity’ slightly improved the map). Automatically sharpened maps from CryoSPARC’s Non-uniform refinement were used in ChimeraX (Goddard et al. 2018) to make figures.

Image processing of 601-DNA nucleosomes bound to Dsup was performed in CryoSPARC. A 2D classification-curated particle stack was extracted at the original pixel size (0.935 Å) and refined using an initial 3D reference with one DNA end wrapped around the octamer (Supplemental Fig. S1 and Supplemental Movie S3). It should be noted that starting the 3D refinement with a 3D reference lacking both DNA ends poorly handles 2-fold averaging around the dyad axis, misrepresenting nucleosome breathing in crosslinked Dsup-bound nucleosomes during subsequent 3D variability analysis (Supplemental Movie S4). After refinement, 3D classification into 6 classes, without alignment, yielded one class whose particles were refined to reconstruct Dsup-bound nucleosome structure I. Particles from 4 other classes were joined and refined to reconstruct Dsup-bound nucleosome structure II.

### 3D variability analysis

After 3D classification with alignment in RELION, selected particles of the native 167-bp 5S rDNA nucleosome were imported to CryoSPARC and a non-uniform refinement run. 3D variability analysis (Punjani and Fleet 2021) was performed with the particles oriented as in the previous refinement, applying a 3D soft mask around the nucleosome and solving three orthogonal principal modes. Results were displayed using ‘Output mode - intermediates’, downsampling box size from 208 to 104 pixels and low-pass filtering the maps at 5 Å. All remaining parameters were left with the default values. Frames correspondent to the first (main) and second component of variability were used to record movies in ChimeraX (Goddard et al. 2018) (Supplemental Movies S1 and S2).

To visualize the effect of the initial reference in 3D refinements (density absent for one DNA end versus density absent for the two DNA ends) on 2-fold averaging of Dsup-bound nucleosomes, 3D variability analysis was performed as described and movies recorded using the main component of variability (Supplemental Movies S3 and S4).

### Cryo-EM: determination of the nucleosomal DNA orientation

For pdb 3LZ0, 25-bp centered on the dyad (DNA position 0) were mutated to the corresponding sequence of the 5S rDNA using ChimeraX (Goddard et al. 2018). For each cryo-EM map whose DNA orientation we analyzed, this coordinates file was rigid-fitted into the map in the two possible orientations related through a 180° rotation over the dyad axis. Separately, each orientation was real-space refined in Coot 0.9.4 (Emsley et al. 2010) followed of Phenix 1.20 (Afonine et al. 2018). Refined DNAs were opened in Chimera 1.15 (Pettersen et al. 2004) to obtain a list of atomic B-factors (Urzhumtsev et al. 2022), excluding the P, OP1, OP2, O5’ and O3’ atoms. These phosphate backbone atoms were excluded because they are subject to greater variations in the value of the B-factors (for instance, the phosphate backbone facing the octamer has in principle much lower B-factors than the atoms facing the solvent). However, in our experience, excluding the P, OP1, OP2, O5’ and O3’ atoms should not significantly alter the final B-factor analysis. Using Microsoft Excel (https://office.microsoft.com/excel), atomic B-factors were averaged per-bp and plotted versus the bp number. Note that to compare the two DNA orientations one was flipped horizontally in the graphic (this meaning that bp −12 to +12 of a given orientation need to overlap with bp +12 to −12 of the opposite orientation).

### Cryo-EM: atomic modelling

In Chimera 1.15 (Pettersen et al. 2004), the nucleotides and amino acids of pdb 7LYA were mutated to match our DNA and histone sequences. Chains were renumbered. The resulting coordinates file was rigid-fitted in the DNA orientation previously found. If needed, DNA segments without EM density in the maps filtered according to local resolution estimation were deleted. Each nucleotide and amino acid was fitted into the density using Coot 0.9.4 (Emsley et al. 2010). When applicable, some amino acids or side chains were removed or de novo built. Real space refinement in Phenix 1.20 (Afonine et al. 2018) was run. 2 to 3 cycles of Coot 0.9.4 and Phenix 1.20 were performed.

